# TWO-SIGMA: a novel TWO-component SInGle cell Model-based Association method for single-cell RNA-seq data

**DOI:** 10.1101/709238

**Authors:** Eric Van Buren, Ming Hu, Chen Weng, Fulai Jin, Yan Li, Di Wu, Yun Li

## Abstract

In this paper, we develop TWO-SIGMA, a TWO-component SInGle cell Model-based Association method for differential expression (DE) analyses in single-cell RNA-seq (scRNA-seq) data. The first component models the probability of “drop-out” with a mixed-effects logistic regression model and the second component models the (conditional) mean expression with a mixed-effects negative binomial regression model. TWO-SIGMA is extremely flexible in that it: (i) does not require a log-transformation of the outcome, (ii) allows for overdispersed and zero-inflated counts, (iii) accommodates a correlation structure between cells from the same biological sample via random effect terms, (iv) can analyze unbalanced designs (in which the number of cells does not need to be identical for all samples), (v) can control for additional sample-level and cell-level covariates including batch effects, (vi) provides interpretable effect size estimates, and (vii) enables general tests of DE beyond two-group comparisons. To our knowledge, TWO-SIGMA is the only method for analyzing scRNA-seq data that can simultaneously accomplish each of these features. Simulations studies show that TWO-SIGMA outperforms alternative regression-based approaches in both type-I error control and power enhancement when the data contains even moderate within-sample correlation. A real data analysis using pancreas islet single-cells exhibits the flexibility of TWO-SIGMA and demonstrates that incorrectly failing to include random effect terms can have dramatic impacts on scientific conclusions. TWO-SIGMA is implemented in the R package `twosigma` available at https://github.com/edvanburen/twosigma.

## 1 Introduction

Advancements in single-cell sequencing technologies have created many exciting opportunities to researchers yet have also posed many challenges relating to data analysis. Expression profiles can now be analyzed at the single-cell level, providing new insights into the cellular heterogeneity of gene expression. Three characteristic features of single-cell transcriptome sequencing data include excess zero counts, overdispersion of observed counts, and a large number of cells that are sequenced from a relatively small number of samples [29]. Technological limitations including low capture rate and amplification failure lead to “drop-out” events, in which the data may capture only a fraction of the transcriptome of a given cell and mistakenly generate zero measurements for expressed genes. The presence of these zeros creates a dataset with an excess of zeros (often called “zero-inflation”) beyond those that occur due to biological factors; these excess zeros often necessitate special modelling approaches such as a two-component model [5, 9]. Overdispersion, in which the variance in expression exceeds the mean expression, is commonly observed in count-based quantitative sequencing due to large variance in expression and within gene variability over time or across samples. A within-sample correlation is also present because multiple cells are sequenced from the same biological sample. These challenges motivated us to develop the new statistical method TWO-SIGMA. It is designed for association analyses where the primary interest is in performing statistical inference on covariate(s) of interest, such as a treatment effect. As we will discuss, TWO-SIGMA is not limited to a two-group comparison and can test for more general kinds of differential expression (DE) while simultaneously controlling for multiple sample-level and cell-level covariates and accounting for within-sample correlation.

Most existing methods for DE analysis in scRNA-seq data are designed for a two-group comparison. As a result, benchmarking papers typically limit themselves to two-group comparisons [28, 23]. Three of the most popular methods for two-group comparisons are SCDE, scDD, and DESingle. SCDE and scDD are both innovative Bayesian methods, with the former utilizing a two-component negative binomial mixture method and the latter using a Dirichlet mixture process [10, 11]. Although both methods show strong performance, only the latter can adjust for confounding covariates, and this adjustment is indirect through a residualized analysis. DESingle employs a zero-inflated negative binomial (ZINB) distribution to analyze DE in scRNA-seq data while accounting for excess zeros and overdispersion [18]. Like SCDE and scDD, however, DESingle does not employ a regression modeling framework to control for other covariates or account for within-sample correlation.

MAST was introduced as a hurdle model for the analysis of scRNA-seq data [5], and is considered to be one of the preferred methods for performing DE analysis in scRNA-seq data [15]. Like TWO-SIGMA, but unlike the methods described above, MAST can test for DE in cases beyond a two-group comparison. There are several important differences, however, between TWO-SIGMA and MAST. First, we fit a zero-inflated model on the observed counts while MAST fits a hurdle model (described in more detail in the next section) on the log scale. The ability to avoid log-transforming the data is desirable given recent evidence which suggests that log transformation can distort many scRNA-seq datasets by producing false variability [25, 16]. Further recommendations for DE analysis in scRNA-seq data state that the observed counts should be modeled directly while accounting for batch effects as covariates rather than through normalization [15]. Second, TWO-SIGMA allows the covariates in each of the two components to differ. We will discuss reasons that this flexibility can be appreciated by researchers later. Third, and most importantly, although the ability to include random effects in either component of MAST is mentioned by its authors, they do not prioritize their inclusion for scRNA-seq data and do not evaluate the impact of random effects on the model’s performance. We will revisit the comparison with MAST in the methods section.

Several unsupervised methods, such as ZINB-WaVE [22] and ZIFA [20] have also been proposed for scRNA-seq analysis. Both ZINB-WaVE and ZIFA are primarily designed for settings in which dimension reduction, not association analysis, is the primary goal. One interesting use of ZINB-WaVE is to construct observation-level weights that can be incorporated into the popular bulk RNA-seq pipelines found in the `DESeq2` [14] or `edgeR` [17] Bioconductor packages [26, 27]. These pipelines do not allow for random effects or model excess zeros separately and can involve some transformation of the data in processing or analysis.

The two-component zero-inflated model without random effects has a long history in the analysis of count and microbiome data [12, 3, 7], however its application in scRNA-seq data is limited. Furthermore, a zero-inflated negative binomial mixed effects model has previously been proposed for modelling zero-inflated count data [19]. The focus of that work was on typical repeated measures applications where the number of repeated measures per individual is small, in contrast to genomic applications which tend to include more repeated measures than samples. Because attention is split between zero-inflated and a similar approach called a hurdle model, and between the Poisson and negative binomial distributions, details regarding the performance and robustness of the zero-inflated negative binomial mixed-effects model are not discussed in as much detail as we can here.

The rest of the article proceeds as follows: first, we specify the TWO-SIGMA model, discuss implications of its parameterization, and provide details on parameter estimation. Next, we describe both traditional methods and a new *ad hoc* method to decide whether random effects should be included in our zero-inflated negative binomial model. Then we show simulation results and an application to a dataset of pancreatic islet single-cells, respectively. Finally, we conclude with a discussion.

## 2 Materials and Methods

### 2.1 Zero-inflated negative binomial distribution

For a given gene, let *i* index the samples sequenced and *j* index the *n*_*i*_ single cells from sample *i*. Consider the following parameterization of the negative binomial probability mass function (p.m.f.) at a non-negative integer *y*_*ij*_ corresponding to the observed read count:

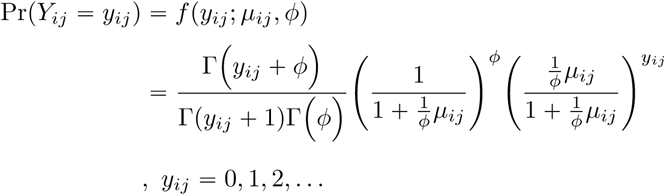

With this parameterization, E(*Y*_*ij*_) = *µ*_*ij*_ and

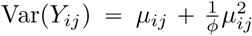, such that *ϕ* is the overdispersion parameter (*ϕ* > 0). This parameterization is appealing for interpretability because as 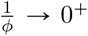, the density above approaches the Poisson density with mean *µ*. Thus, as mentioned further in the discussion, the Poisson and negative binomial distributions are asymptotically nested (and nearly identical for large values of *ϕ*).

To accommodate the excess zeros often observed in scRNA-seq data, we employ the zero-inflated negative binomial distribution (ZINB). This distribution mixes a point mass at zero (from which observations are considered “drop-out”) with the negative binomial distribution. Let *p*_*ij*_ and *µ*_*ij*_ be the probability of drop-out and the mean read count conditional on not being dropped-out for cell *j* from sample *i*, respectively. The p.m.f. of one observation *Y*_*ij*_ under the ZINB distribution is given by:

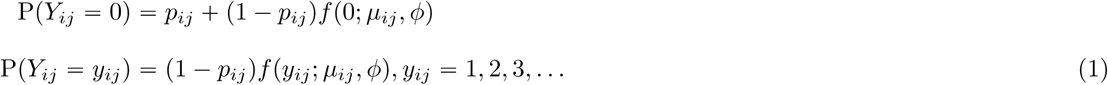

The ZINB distribution thus assumes that there are two sources of zeros in the data: the first source is the process that governs drop-out and the second source is from the negative binomial process for genes that are not dropped-out. This differs from hurdle models used for gene expression data [5] which use a left-truncated or continuous distribution for the positive expression component—meaning zeros can only be generated from the drop-out process. Thus, the hurdle model does not allow zero expression measurements due to biological variation.

Interpretations from the zero-inflated model are quite natural for single-cell gene expression data because it is reasonable to believe that some observed zeros are “structural zeros” with *bona fide* zero expression due to stochastic biological factors (e.g. transcriptional bursting, cell cycle) and not due to technical drop-out [29]. Although semantic, the distinction regarding the source of zeros affects the interpretation of model coefficients and is important because these models are often misinterpreted by researchers [21, 24].

### 2.2 TWO-SIGMA

We can now provide the full TWO-SIGMA specification:

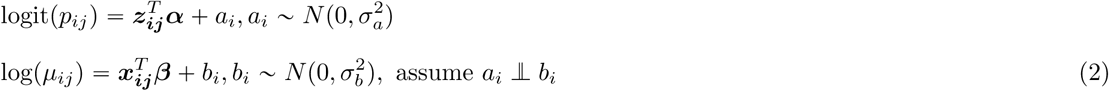

The model is fit for each gene individually, so all parameters are gene-specific. ***α*** and ***β*** are fixed effect coefficient vectors and the corresponding vectors of covariates ***z***_***ij***_ and ***x***_***ij***_ can be different. *a*_*i*_ and *b*_*i*_ are sample-specific intercepts (discussed more in the next section). Prediction of sample-specific intercepts and estimation of the variance components 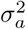 and 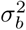 allow us to investigate heterogeneity among individuals, and tests of whether the variance components equal zero allow us to separately (or jointly) evaluate the need for random effects. Separate variance components are estimated because the different link functions in the two components correspond to linear predictors with different scales. Including the random effects terms also helps control for any within-sample correlation, providing more accurate estimates and standard errors of fixed effect parameters.

As part of our `twosigma R` package, we employ the `glmmTMB` package [2] to fit the model specified in equation (2). This package is well-suited to fit generalized linear mixed models (GLMMs) because the user can easily specify an arbitrarily complex model composed of fixed and random effects. More details regarding computational considerations can be found in section 5 of the supplement. To summarize, TWO-SIGMA controls for additional covariates in both components, incorporates random effects to accommodate within-sample dependency, can analyze unbalanced data, and allows for zero-inflated and overdispersed counts. The regression modelling framework provides interpretable effect size estimates and can examine any DE hypothesis that can be expressed as a contrast of regression coefficients. The implementation of the model strikes a balance between computational accuracy and efficiency, even as the number of random effects (number of samples in the context of the scRNA-seq data) or the number of single cells per sample increases.

### 2.3 Evaluating the need for random effects

One primary methodological contribution of TWO-SIGMA for scRNA-seq data analysis is the inclusion of random effect terms in each of the two components, which is a well-established technique to account for within-sample correlation. Ignoring random effects in TWO-SIGMA is equivalent to assuming that cells from the same sample/individual are independent. This assumption can lead to underestimated standard errors and thus inflated type-I errors. An example is given in table 3 in the real data analysis section.

**Table 1:**
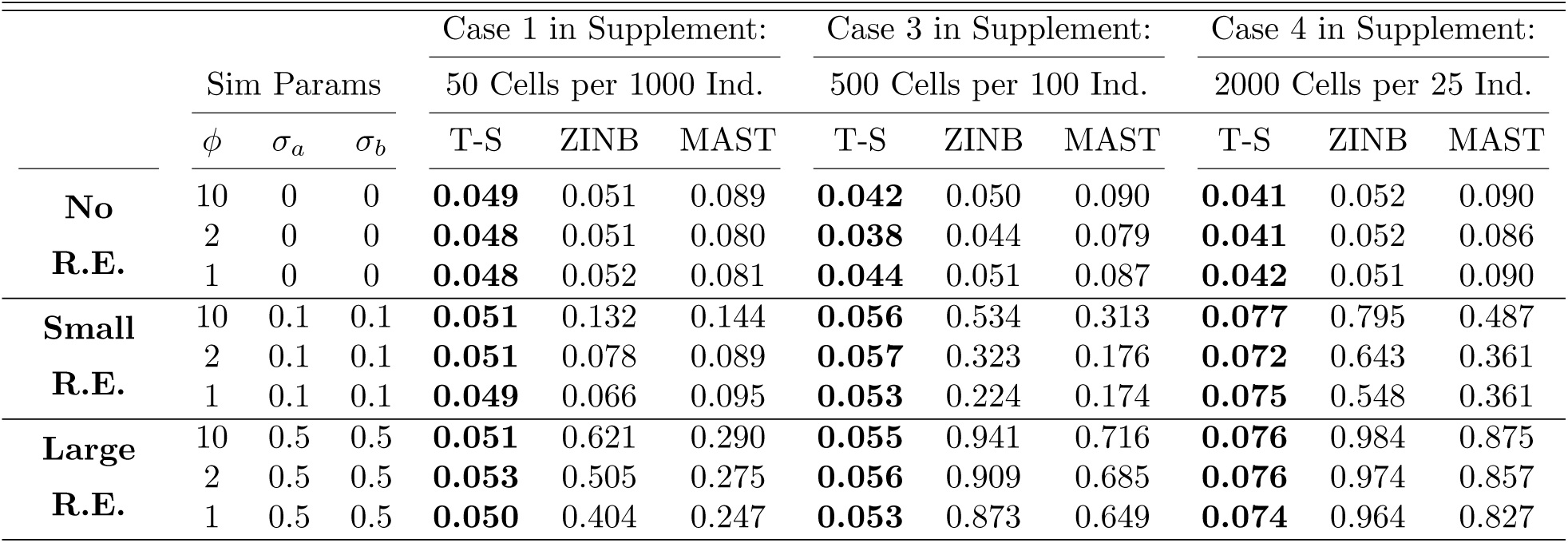
Type-I error evaluations in simulated data: Shows type-I error using the LRT to test the joint null hypothesis of a simulated binary disease status indicator, H_0_ : *α*_1_ = 0, *β*_1_ = 0 versus H_*a*_ : *α*_1_ ≠ 0 or *β*_1_ ≠ 0, with a significance level of 0.05. “T-S” refers to TWO-SIGMA, ZINB refers to a zero-inflated negative binomial model without random effects and MAST refers to the model described in [5]. 10,000 genes were simulated.

**Table 2:** Rejection summaries from the pancreas data: Shows the proportion of genes in the pancreatic islet data with rejected nulls for various hypotheses related to T2D. The TWO-SIGMA model as specified in equation (2) was fit with no zero-inflation variance component (no ZIVC).

| Hypothesis | Alpha Cells | Beta Cells |
| --- | --- | --- |
|  | No ZIVC | No ZIVC |
| Overall Disease Status | 0.161 | 0.111 |
| Overall R.E. Test | 0.738 | 0.724 |
| NB vs. Poisson | 0.627 | 0.555 |

**Table 3:** Influence of failing to include needed random effects: Gives mean component estimates for gene *RPS29* with (top panel) and without (bottom panel) random effects

| Effect | Estimate | Std. Error | z value | p-value |
| --- | --- | --- | --- | --- |
| Intercept | 0.521 | 0.207 | 2.515 | 0.012 |
| T2D | -0.349 | 0.292 | -1.197 | 0.231 |
| age | -0.284 | 0.256 | -1.109 | 0.267 |
| CDR | 0.394 | 0.011 | 36.284 | <.001 |
| $\sigma_b$ | 0.490 | | | |
| Intercept | 0.833 | 0.021 | 40.094 | <.001 |
| T2D | -0.605 | 0.032 | -19.090 | <.001 |
| age | -0.057 | 0.017 | -3.324 | <.001 |
| CDR | 0.390 | 0.015 | 26.611 | <.001 |

Evaluating the need for a random effect term involves a hypothesis test of whether the corresponding variance component(s) equal zero. For example, consider testing whether random effects are needed in either component of TWO-SIGMA with H_0_ : *σ*_*a*_ = *σ*_*b*_ = 0 versus H_*a*_ : *σ*_*a*_ > 0 or *σ*_*b*_ > 0 using the Likelihood Ratio Test (LRT). This procedure requires fitting the model under both the null and the alternative hypotheses, but is a preferred method to determine whether the random effect terms significantly improve model fit. Other less desirable post-fitting options to compare models with and without random effects include information criteria like AIC and BIC or Wald tests of the variance components [6]. Critically, all options discussed require fitting the “full” model including the random effect terms. The scRNA-seq application is distinct from typical repeated measures analyses in that the number of repeated measures (cells) typically far exceeds the number of samples. Such designs can entail more extensive computational time for each gene over scenarios involving a smaller number of repeated measures from a modest number of individuals. These computational burdens are especially relevant given that scRNA-seq data typically include thousands of genes, each of which is fit separately using TWO-SIGMA. It would therefore be useful to identify the genes that are most likely to need the random effect terms without having to fit the full model to each gene.

#### 2.3.1 ad hoc approach

We utilize the following *ad hoc* approach to determine whether random effects are needed: using a one-way ANOVA, we regress the Pearson residuals from a zero-inflated negative binomial regression model without random effects on the sample label and take the *p*-value from the overall ANOVA *F* test. This *p*-value serves as a rudimentary measure of whether the residuals tend to differ across samples. If they do, this is evidence that residuals are not exchangeable across samples. The full TWO-SIGMA specification including random effects will then be fit to more formally evaluate the need for random effect terms. In contrast, when the residuals show no tendency of differing across samples, we do not have evidence to believe that they are structured/clustered within samples and thus will not fit the full model with random effects. Through simulations we found that this procedure is very effective in identifying the need for random effects. Results from applying this proposed method to a real dataset of pancreatic islet cells are given in the data analysis section. In simulations, computation runtime was the longest for models attempting to fit random effects when variance components were truly zero (see supplementary tables 1-4). Therefore, as discussed more in section 3 of the supplementary file, the *ad hoc* method can dramatically reduce overall computation time over many genes in addition to increasing model parsimony where most appropriate.

### 2.4 Simulation studies

To evaluate the performance of TWO-SIGMA, we simulated data in a variety of scenarios. Although many methods exist for DE in scRNA-seq data, as described above, we chose to focus our comparison to MAST because, like TWO-SIGMA, it uses a regression modeling framework that is suitable for designs beyond a two-group comparison and can simultaneously control for multiple cell-level and subject-specific covariates. We also compare to a ZINB model without random effects to highlight the impact random effect terms can have on model performance. Simulated covariates included disease status, age and the cellular detection rate (CDR, see the real data analysis section and [5] for more details). Values of ***α*** and ***β*** were designed to mimic realistic parameter values observed in our pancreatic data analysis. Models were evaluated using the likelihood ratio test on the joint null hypothesis that a disease status indicator is not associated with expression through either drop-out probability or the conditional mean, H_0_ : *α*_1_ = *β*_1_ = 0. We consider two different ways of simulating data: one in which the number of samples far exceeds the number of cells per sample, as is typical in most repeated measures contexts, and the other in which the number of cells far exceeds the number of samples, as is the case in scRNA-seq data. In each scenario we simulated 10,000 genes and used 0.05 as the nominal significance rate to evaluate type-I error and power.

## 3 Results

### 3.1 Type-I error control

Table 1 shows results from simulations in which the true values of the overdispersion parameter *ϕ* and the variance components *σ*_*a*_, *σ*_*b*_ vary. Type-I error is well-controlled for TWO-SIGMA in the scenarios involving more individuals than cells. When the number of cells increases, type-I errors from TWO-SIGMA are slightly inflated over the nominal rate of 5%, but consistently remain superior to the results from the ZINB model or MAST in the presence of ignored non-zero variance components. For example, the last row of table 1 shows that, when *ϕ* = 1 and *σ*_*a*_ = *σ*_*b*_ = 0.5, type-I error for TWO-SIGMA increases from 0.05 to 0.053 to 0.074 as the number of individuals decreases from 1000 to 100 to 25. In contrast, the ZINB model and MAST have inflated type-I errors in every scenario that increase to nearly 1 as the number of individuals decreases. This is not surprising because both of the latter methods cannot account for any within-sample dependency structure among the single cells from the same sample. Ignoring the dependency introduced by even a moderate random effect size can thus have a drastic impact on the type-I error. When true variance components are zero, both TWO-SIGMA and the ZINB model preserve type-I error while MAST consistently has higher type-I error, as seen in the first three rows of table 1. Coverage of confidence intervals for *α, β*, and *ϕ* always approaches the nominal level (Supplementary tables S1-S4). The reason for the slightly inflated type-I error for TWO-SIGMA observed in the scenario with 25 individuals is worth mentioning briefly. The smaller number of individuals (25) provides less information to estimate the sample-specific variance components *σ*_*a*_ and *σ*_*b*_ and few unique values of the simulated binary disease status indicator. The slightly lower coverage for variance components in the last 6 sets of supplementary table 4 is one illustration of the (relative) difficulty in getting precise variance component estimates. TWO-SIGMA outperforms MAST or the ZINB model in preserving type-I error and estimating parameters under a variety of sample size breakdowns and with a variety of true parameter values. See supplementary figures S4-S5 for type-I error across more stringent significance thresholds for representative scenarios.

### 3.2 TWO-SIGMA retains high power under a variety of scenarios

Because the ZINB model and MAST both have heavily inflated type-I errors in many cases, using raw (or “apparent”) power does not provide a fair comparison for these two methods. For each method and each simulation setting under the null, we therefore calculate the empirical significance threshold, defined as the test statistic value at the quantile associated with 1 minus the significance level. A percentage of statistics equal to the nominal significance level will then be larger than this threshold. For various alternative hypotheses, we calculate “true” power for MAST and the ZINB model by using the empirical significance threshold from the corresponding setting under the null as the rejection threshold instead of a usual theoretical threshold (e.g. 5.9915 from 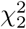 at the .05 level). Figure 1 plots raw power for TWO-SIGMA and true power for MAST in the ZINB model in the following four scenarios: effect in both components, in either the same or opposite directions, and effects in only one of the two components. In the first three scenarios, MAST consistently has the lowest power, while TWO-SIGMA and the ZINB model have very similar power in the first two scenarios, beginning at around 20% and increasing to nearly 100%. The ZINB model has higher power than TWO-SIGMA in the third scenario but the lowest power in the fourth scenario. In simulation, computing the empirical significance thresholds and true power is straightforward and computationally included. In real data settings, however, computationally intensive resampling approaches are needed for reliable estimates of the empirical significance thresholds. Because TWO-SIGMA preserves type-I error, we can rely on raw power and can therefore bypass the need for any resampling approach for valid inference. This is a key advantage and shows that TWO-SIGMA is more robust and flexible than the ZINB model while both preserving the type-I error and having high power without any additional computation. When the effect is only in the zero-inflation component, power is lower for all methods than in the first three scenarios. Such effects present only in the zero-inflation component are known to be more difficult to detect, as seen in [3]. For full power results, including more detailed comparisons to the ZINB model with additional discussion, see section 4, figures S6-S8 and tables S5-S12 of the supplement.

**Figure 1:**
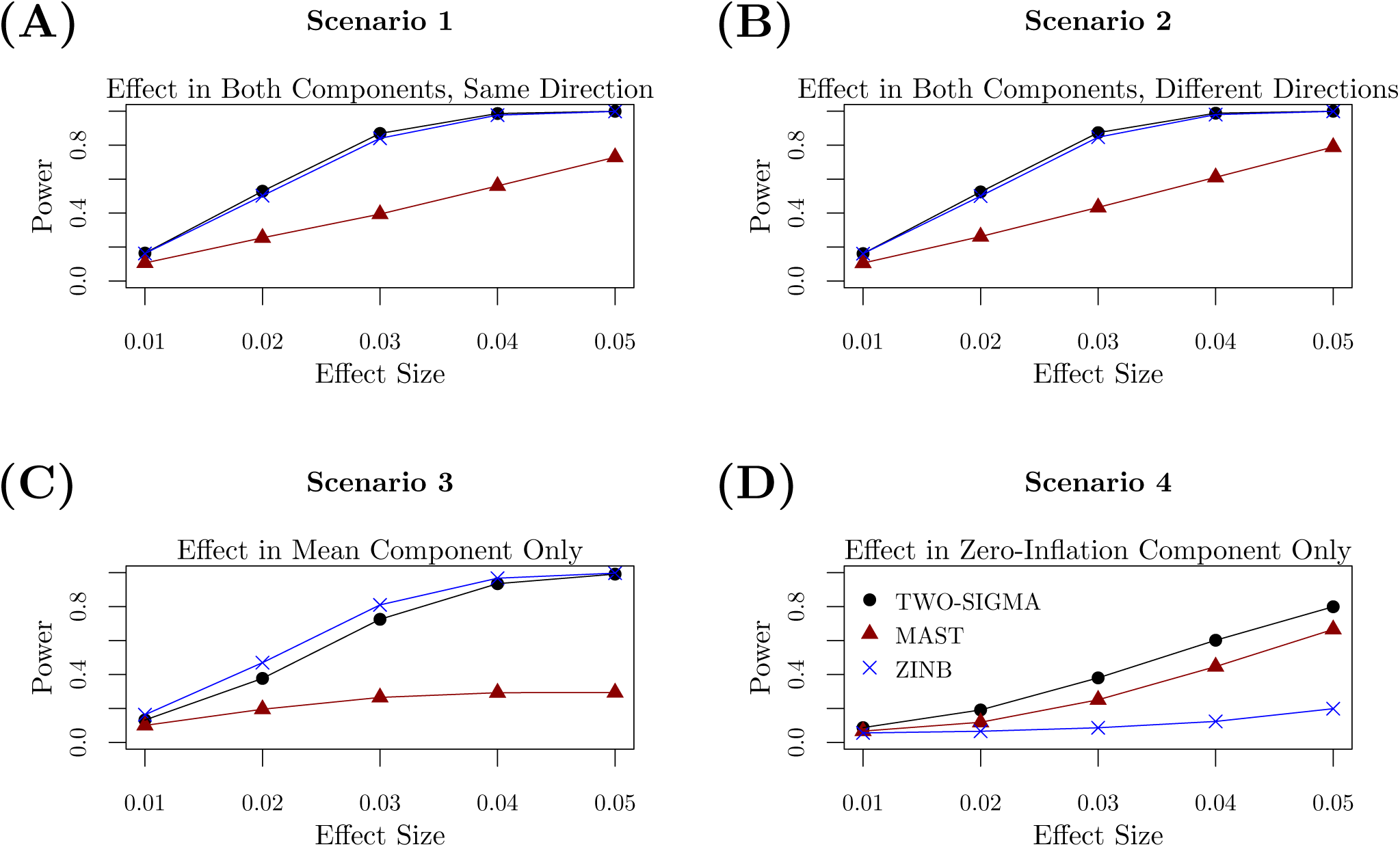
Power evaluations in simulated data: Shows the power to test H_0_ : *α*_1_ = *β*_1_ = 0 by varying the effect size with 500 cells from each of 100 individuals. Values of *ϕ, σ*_*a*_, and *σ*_*b*_ were all set to 0.1 to mimic the “Small R.E.” section of table 1 and 10,000 genes were simulated. Because of the type-I error inflation from the ZINB model and MAST seen in table 1, true power was calculated and plotted for these methods using the empirical significance threshold from the corresponding setting under the null. TWO-SIGMA can bypass the need for computationally expensive resampling procedures needed to generate true power because it preserves the type-I error as seen in table 1. See the discussion at the beginning of section 4 of the supplement for more details about computing true power and discussion regarding power trends across the different methods.

### 3.3 Sensitivity of the *ad hoc* method for random effect screening in simulated data

We evaluated the *ad hoc* screening procedure used to select genes possibly in need of random effect terms using the simulated data. We found it to be effective as a screening procedure: for the data used in Table 1, most genes with non-zero variance components had p-values less than 0.05 and were flagged as in need of estimating variance components using TWO-SIGMA. When the variance components are zero, however, the p-values from the ad hoc method are close to uniformly distributed, indicating that the procedure is not too liberal under the null. See section 2 and figures S2 and S3 of the supplement for more details and some example results.

### 3.4 Pancreas real data analysis

For illustrative purposes we applied TWO-SIGMA to a dataset of pancreatic islet cells isolated from nine individuals (see section 1 of the supplement and [4] for full details on the data processing and generation steps). To focus on the most informative cells and genes, we applied rather aggressive filtering of the data to keep the top 2,000 genes by number of transcripts observed and only keep cells with more than 1000 transcripts across these genes. After merging across all nine individuals, we were left with 1,290 genes and 10,269 single cells of which we used only the 7,774 for which cell type information was available based on the expression of signature genes. Here we focus our attention on alpha and beta cells, which compose the majority (55% and 34%, respectively) of the cells in our dataset. Type-II diabetes (T2D) status is of primary interest, and age is included as an additional subject-level covariate given its potential to confound the relationship between T2D status and gene expression. The cellular detection rate (CDR) is defined in [5] as the percentage of genes expressed over some background level of expression (often chosen to be zero). The CDR therefore has a biological interpretation as a cellular scaling factor and is a surrogate for both technical and biological variation. This confirms the conclusions of [8] that the CDR can explain a substantial proportion of observed expression variability and should be included in any association analysis of scRNA-seq data. As such, we include CDR in all analyses performed and stratified by cell type. For more details about the pancreas data processing, see section 1 and figure S1 of the supplementary file.

Figure 2 plots the relationship of mean versus variance for the 1,290 genes we used in our analysis. It shows that the Poisson and zero-inflated Poisson models cannot adequately account for the overdispersion observed in many genes. In contrast, TWO-SIGMA can accommodate these mean-variance pairs in a quadratic relationship via the overdispersion parameter *ϕ*. Because we have only nine individuals, we chose to focus on analyses excluding the zero-inflation random effect terms *a*_*i*_ to improve convergence and overall model fit. Some genes still showed convergence issues–partly indicative of a misspecified or overparameterized model and partly due to the small number of cells and samples in the dataset. As a general guideline, we recommend that users with concerns or limited computational resources begin including random effects in the mean component, and scale upwards to include random effects in the zero-inflation component if performance is satisfactory.

**Figure 2:**
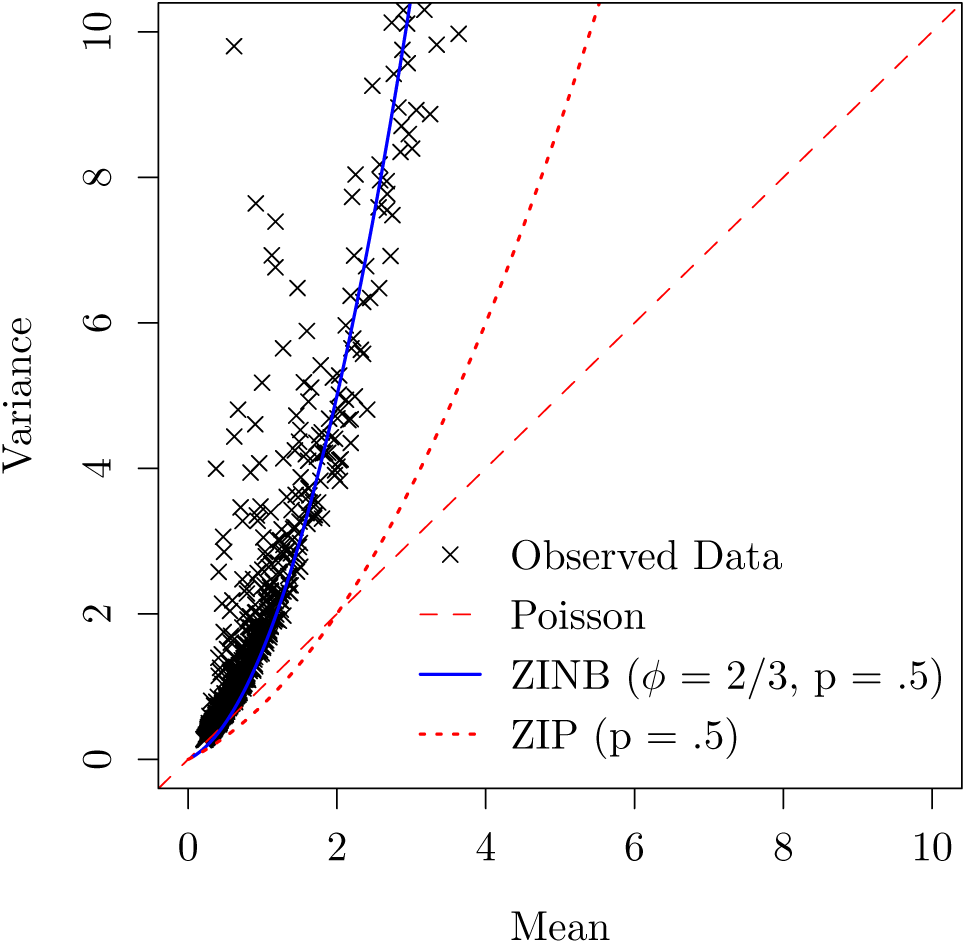
Presence of overdispersion in real data: Shows the need of a non-linear mean-variance relationship in the pancreatic islet data. Each point represents the mean-variance relationship for one gene. In the legend *ϕ* represents the overdispersion parameter of the negative binomial distribution and p represents the drop-out probability.

Table 2 shows the proportions of genes showing statistically significant results at the .05 level for three types of hypothesis tests: the joint test of significance for the binary disease indicator H_0_ : *α*_1_ = *β*_1_ = 0, the test of the mean model variance component H_0_ : *σ*_*b*_ = 0, and the test for the presence of overdispersion 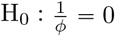. For example, when fitting the TWO-SIGMA model without the zero-inflation variance component to alpha cells, 73.8% of genes had statistically significant variance components in the mean model. Most genes showed the need for a random effect term or the negative binomial distribution (or both).

#### 3.4.1 Impact of ignoring within-sample correlation

Models for genes that mistakenly exclude the *b*_*i*_ random effect term often show highly significant results for covariates; this significance can disappear when including the random intercept term—possibly indicative of a false positive due to failing to account for within-sample correlation. For example, gene *RPS29* demonstrates this pattern in alpha cells. Table 3 shows that failing to include random effects—and thereby assuming independence of all single cells—can lead to vastly underestimated standard errors. T2D status and age change from highly significant to insignificant when including a random intercept term. The standard error for the coefficient of T2D increases by a factor of 9 from 0.032 to 0.292, and the magnitude of the point estimate is halved from −0.605 to −0.349. Individual covariates such as T2D can thus exhibit dramatically increased type-I error when random effects are incorrectly ignored. In contrast, the coefficient and associated standard error for the cellular detection rate (CDR) are nearly identical in the two models. This result is expected given that CDR is a cell-level covariate and shows that including sample-specific random effects leads to very minor changes in the estimation of any covariates that are not sample-specific. Our emphasis in this section is not to draw conclusions about any association between *RPS29* and T2D, but rather to illustrate that ignoring random effects has the potential to alter scientific conclusions.

#### 3.4.2 Cell-type specific genes often show a need for random effect inclusion

We matched 234 and 120 genes in our data that were identified in previous studies as cell-type specific in alpha or beta cells, respectively. ([13], supplementary table 10). After stratifying the data by cell type and removing genes with more than 90% or less than 10% zeros, we fit TWO-SIGMA (excluding *a*_*i*_ as mentioned previously) to the remaining 222 alpha cell-specific and 111 beta cell-specific genes to alpha cells and beta cells, respectively. Of these, 93 alpha cell-specific genes and 85 beta cell-specific genes had statistically significant variance components *σ*_*b*_. This suggests that non-negligible between-sample variation—not attributable to cell-type—is present for these cell-type specific genes. As discussed in [13], cell-type specific expression profiles are often of primary interest to study (dis)function at the cellular level and reveal novel approaches to treat and manage diseases such as T2D. Thus, it is critical to have reliable inference for these genes. As seen in the previous section, incorrectly excluding random effect terms can provide very misleading results and can thereby misdirect attempts to understand disease etiology at a cellular level.

We also used alpha cells to test the overall effect of T2D using both TWO-SIGMA to MAST. Table 4 shows that MAST rejects in many more instances than TWO-SIGMA. Of the 273 genes that were rejected with MAST but not with TWO-SIGMA, 234 have statistically significant variance components in TWO-SIGMA. This further illustrates the possibility that fixed effect coefficients can be mistakenly deemed significant in the presence of within-sample correlation.

**Table 4:** Agreement between TWO-SIGMA and MAST: Shows the agreement in rejecting the omnibus null hypothesis of an association between T2D status and gene expression in alpha cells using a Bonferroni adjusted significance level of 5×10^−5^.

| TWO-SIGMA | MAST |  |
| --- | --- | --- |
|  | No Reject | Reject |
| No Reject | 1013 | 273 |
| Reject | 1 | 3 |

#### 3.4.3 The *ad hoc* method successfully separates genes that need random effects

Finally, we used all 1,290 genes from the islet dataset to demonstrate the usefulness of the *ad hoc* method to determine the need for the random effects terms *b*_*i*_. Figure 3 shows that likelihood ratio statistics from formal testing of *b*_*i*_ are consistently larger for genes selected by the procedure than those not selected. This pattern suggests that the *ad hoc* procedure described earlier can effectively identify genes that will exhibit non-zero variance components in real data.

**Figure 3:**
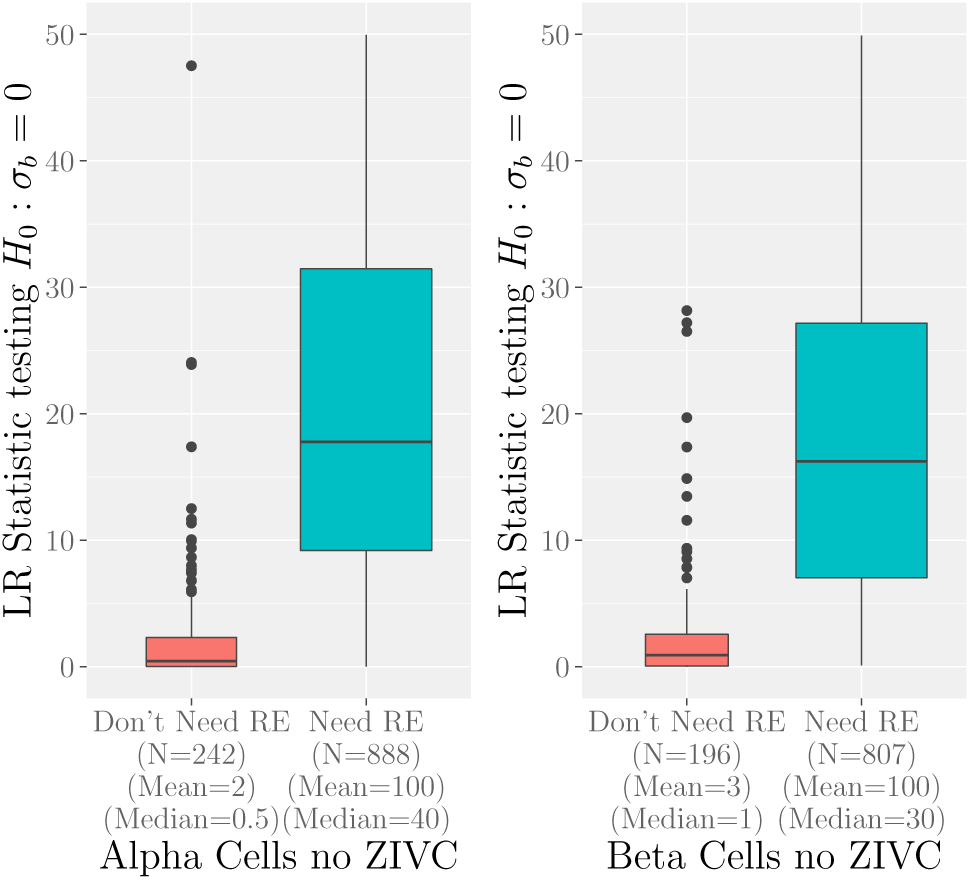
Ability of the ad hoc method to identify genes in need of random effects: Shows boxplots of the LR statistics from the joint test of the need for random effects, *H*_0_ : *σ*_*a*_ = *σ*_*b*_ = 0, using TWO-SIGMA. Genes that our *ad hoc* procedure suggests need random effects (“Need RE”) and genes the procedure suggests do not (“Don’t Need RE”) are compared. Both panels were created using TWO-SIGMA as specified in equation (2) but with no zero-inflation variance component (no ZIVC).

## 4 Discussion

We have developed TWO-SIGMA, a two-component zero-inflated negative binomial model with random effects for association analysis of scRNA-seq data. The model builds on the well-established literature in both zero-inflated models and generalized linear mixed models. It keeps the data on the original scale while simultaneously allowing for zero-inflation, overdispersion, and random effects to account for within-sample correlation. As compared to existing methods, its flexibility is demonstrated both in the use of random effect terms and the ability to test any hypothesis of DE that can be expressed as a contrast of regression coefficients while controlling for multiple sample-level and individual-level covariates.

Including random effects explicitly controls for within-sample correlation, and can improve mean parameter and standard error estimates. Given that many scRNA-seq studies have few samples, it would be reasonable to consider controlling for sample as a fixed effect rather than a random effect. However, there are two reasons to prefer incorporating a random effect to a fixed effect approach. First, we are interested in estimating a variance component that can apply to all samples in the population. Second, the random intercepts explicitly control for the within-sample correlation, rather than only providing adjusted parameter estimates for included covariates. An alternative approach to accounting for such sample-level repeated measures would be to fit a marginal model with the generalized estimating equations (GEE) approach instead of a mixed effects model [1]. We chose not to do so for two main reasons: first, we retain the flexibility for sample-level prediction. Second, given that many scRNA-seq experiments are conducted over a small number of samples, it is likely that the empirical (sandwich) covariance estimate would underestimate the true standard errors [1].

Incorrectly excluding random effects and assuming independence of cells can lead to underestimated standard errors of fixed effects and can therefore increase the type-I error of hypothesis tests relating to fixed effects parameters. See table 3 for an example. If the random effect terms do not contribute to the model fit, as judged by a statistical test or practical significance, they can be removed easily within the general framework of TWO-SIGMA. Random intercepts can also be useful even when they are not of direct interest: they often capture the effects of omitted sample-specific covariates, and can limit the bias of fixed effect coefficients caused by misspecification. For example, if cell-type information is missing, and varies between individuals, a random intercept term can limit the resulting bias and p-value inflation observed in fixed effects parameters. Our *ad hoc* method proves to be a useful tool to both select genes that could benefit from including random effect terms and reduce overall computation time by suggesting genes that do not need to be fit including random effect terms.

Because we expect *a priori* that zero-inflation will occur in scRNA-seq, it is beneficial to include a component dedicated to it. The zero-inflation component in TWO-SIGMA is flexible in that it allows for a different set of covariates from the mean model, or no covariates at all. For example, one might be interested in using zero-inflation only to improve mean parameter estimation. In this scenario, a constant probability of drop-out could be assumed via an intercept-only regression model. This would prevent coefficient estimates in the mean model from being overly shrunk towards zero, as would occur if drop-out was not accounted for, but would also limit the total number of parameters estimated and maximize model parsimony. Even if the data are not truly generated from a zero-inflated process, or if drop-out is viewed as a “nuisance,” using the two component model in equation (2) can be a convenient choice to improve model fit and fixed effect parameter estimation. See section 6 of the supplement and supplementary figure S9 for more discussion regarding the zero-inflation component.

Fitting the TWO-SIGMA model also provides a way to choose between the zero-inflated negative binomial and zero-inflated Poisson distributions that may be useful in a standard data analysis; if the overdispersion parameter estimate is small, one can justifiably reduce model complexity and fit a Poisson model. Specifically, one can test 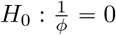 versus 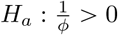 using the likelihood ratio test (p-values come from a 50:50 mixture of 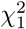 and a point mass at zero). As seen in table 2, the p-value from such a test will often suggest a significant deviation from the Poisson model in scRNA-seq data.

Finally, our experience suggests that variance component estimates are often much smaller in the zero-inflation component than in the mean component. Therefore, as we did in the real data analysis, it might be a pragmatic choice to exclude random effects from the zero-inflation component of TWO-SIGMA. A key strength of TWO-SIGMA is the flexibility to easily customize the model within the general framework either *a priori* or via iterative removal based on statistical hypothesis tests of features such as random effects, overdispersion, or the drop-out component.

## Supporting information

Updated Supplementary File

## 5 Data Availability

The pancreas Drop-seq data is available at GEO: GSE101207.

