## Supplementary material for "TWO-SIGMA: a novel TWO-component SInGle cell Model-based Association method for single-cell RNA-seq data": Updated Supplementary File

### 1 Pancreas Data Methods

#### 1.1 Drop-seq reads processing

We performed raw reads processing following the instructions described in the original Drop-Seq publication [1]. The sequenced Drop-Seq libraries yield 50-base paired-end reads (PE50). However, since only the first 20bp of read 1 is informative (base 1-12 cell barcode, base 13-20 UMI), we trimmed base 21-50 of read 1 before further analysis. We first removed all data with the quality score of read 1 (base 1-20) lower than 10. Read 2 was trimmed at 3' end to remove ployA tails of at least 6 bases, and trimmed at 5' if template switching oligo (TSO) adapter sequence appears. Clean reads were then aligned to hg19 or mm9 using STAR with default settings. We only keep uniquely mapped reads on gene exons. We next filtered out PCR duplicates with the same coordinates, cell barcode, and UMI. We then grouped the reads by cell barcode, and generated the digital UMI-count matrix after counting transcripts for every genes with every cell barcode.

#### 1.2 Distinguish cell barcodes with single cell transcriptomes

We defined STAMPs (single cell transcriptome attached to microparticles) as cell barcodes with significantly more reads than background. Under the Drop-Seq experimental settings, only about 2-5% of beads are co-encapsulated with cells. Therefore, most cell barcodes only have a small number of transcripts from mRNA contamination during the bead breakage step. In order to distinguish STAMPs from empty beads, we examined the density plots of transcript counts for all cell barcodes (see Supplementary Figure 1B of [2]). In all experiments, we observed a major peak from empty beads and a fat right tail representing STAMPs with single cell transcriptomes. We therefore took a simple approach by calculating mean ($\mu$) and standard deviation ($\sigma$) of the major peak assuming a Gaussian distribution. Any cell barcode with more than $\mu + 2 * \sigma$ transcripts were called as STAMPs.

#### 1.3 Down-sampling sequencing data

Because the Drop-Seq libraries have different sequencing depth, we observed variable sensitivity in detecting transcripts / genes from each library (see Supplementary Figure S1), which causes bias during the clustering or comparative analyses. We therefore took a down-sampling approach to normalize the sequencing depth. We firstly run raw data processing as described above using full data and estimate the total STAMP numbers for each donor. For down sampling, we only took a portion of reads from every library so that the average per-STAMP sequencing depth are similar. New UMI-count matrices are generated again for all donors after down sampling. We found that the normalization of sequencing depth

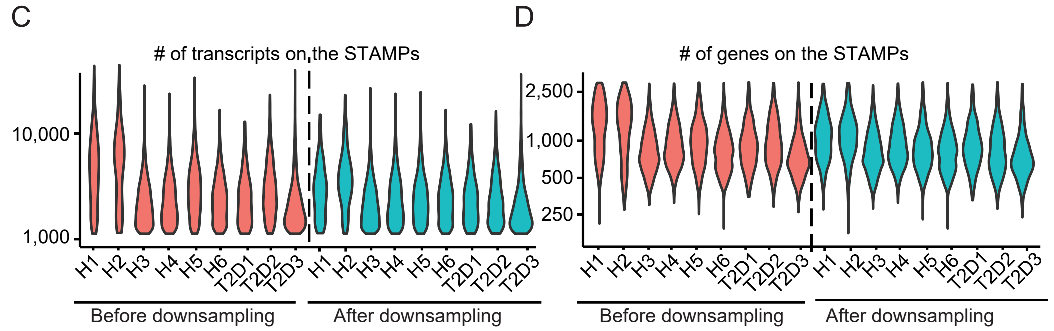


Supplementary Figure S1: Shows the influence of the down-sampling procedure by individual in both the number of transcripts and the number of genes in the STAMPs.

resulted in cleaner clusters in t-SNE plot (see Supplementary Figure 1E of [2]). We only used the down-sampled data matrices when different donors need to be compared.

#### 1.4 Cell type identification using unsupervised clustering

We designed a pipeline to determine the cell types of most STAMPs with high confidence using unsupervised clustering methods (see Supplementary Figure 1A of [2]). Firstly, we performed initial clustering analysis with the 11,920 top STAMPs with at least 1,000 transcripts after down sampling. It has been previously estimated that in human islets, $< 0.1\%$ endocrine cells are positive with more than one marker hormones (INS, GCG, PPY, SST, and GHRL). We therefore first filtered out 890 STAMPs (out of 12,810, or 6.9%) expressing two hormones (see Supplementary Figure 1G of [2]) before clustering analysis. In this step, one STAMP is considered as doublets if it has two hormone genes with 15 transcripts. As mentioned above, the percentage of doublets is significantly greater than estimated from species mix experiment because single cells from tissues are more inclined to adhere with each other than cultured cells.
For clustering, we first ranked top 10,000 genes based on average expression level among all cells; then grouped them into 10 bins with 1,000 genes each. Coefficient of variation (CV) is calculated for every gene within each bin. From every bin, we pick top 50 genes with highest CV as informative genes. All together, we picked 500 informative genes for clustering analysis. We used Seurat package for clustering analysis with default parameters. In Seurat, PCA was performed with the 500 informative genes. Using PC1 to PC10, cells were embedded in a K-nearest neighbor (KNN graph). Smart local moving algorithm (SLM) was applied to group cells into communities. PC1 to PC10 were used as input to visualize cell clusters in two-dimensional t-SNE space. In order to define cell type, we used Seurat FindMarker function to find marker genes of each cell cluster, and defined cell types based on our knowledge and literatures. We performed the first round clustering to classify non-endocrine cells (ductal cells, active PSCs and quiescent PSCs, Figure 1B) and the second round to distinguish the endocrine cell types ($\alpha$, $\beta$, $\delta$, PP cells, Figure 1E). Acinar and $\epsilon$ cells are not distinguishable in t-SNE plot due to scarcity, but can be clearly recognized from PCA plots (See Supplementary Figure 2 of [2]). We also noticed a very small number of

STAMPs (223, or 1.8%) expressing hormone genes inconsistent with their cell type classification (>15 transcripts) (see Supplementary Figure 1H of [2]), which were also filtered as possible doublets after clustering analysis (see Supplementary Figure 1A of [2]). Finally, we successfully assigned unique cell types to 11,697 STAMPs with high confidence. We used the same method for other clustering analyses in this work.

##### 1.5 Cell type identification of low-depth STAMPs

Finally, we classified the low-transcript STAMPs using the knowledge obtained from clustering the top STAMPs as the training dataset (see Supplementary Figure 1A of [2]). As mentioned above, we performed PCA clustering of the training dataset using 500 informative genes. From the PCA results, we took 32 significant principle components (PCs) as knowledge learned from training set. The 32 PCs are linear combinations of the 500 informative genes, and compose a virtual 32-dimensional space. Each cell type should form a cluster in the space. We next calculated the arithmetic centers of 8 cell types from training dataset (ductal, acinar, PSC, $\alpha$, $\beta$, $\delta$, $\epsilon$, PP cells), and built spheres for all cell types centered at their arithmetic mean in the 32-dimensional space. We also computed the Euclidean distance between every cell to the center of its cell type, and empirically defined the radius of each "cell type sphere" as 80 percentile of all the distances in this cell type. For any low depth STAMP, we also took the same 500 informative genes, computed its projection onto the 32 PCs from training data, and the distances between the STAMP and the centers of all cell type spheres. If one STAMP is located exclusively in one cell type's "sphere", we will annotate the STAMP to that cell type. We also performed several filtering steps similar to training set, and successfully classified 16,329 additional STAMPs (see Supplementary Figure 1A of [2]). Lastly, we used 2-dimensional PCA plots and visually confirmed the correctness of cell type assignment (see Supplementary Figure 2 of [2]).

#### 2 More Results from the *ad hoc* procedure

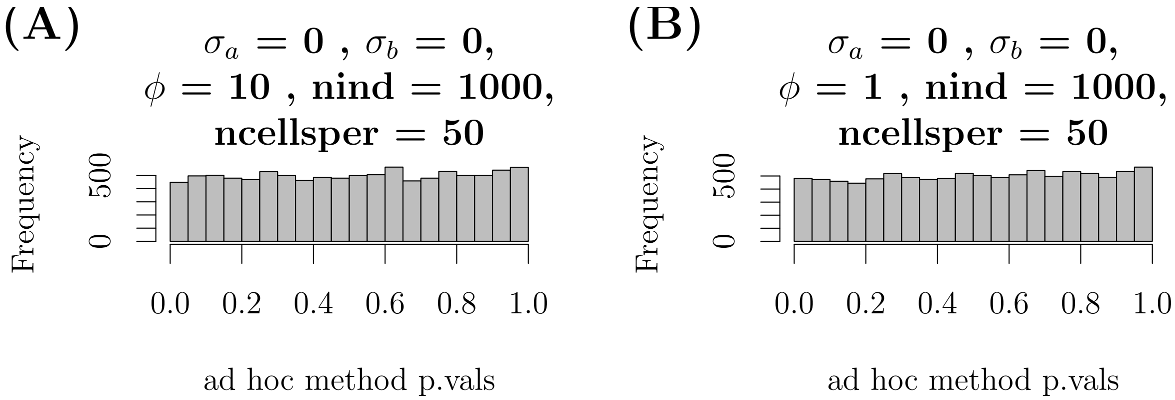


Supplementary Figure S2: *ad hoc* procedure with zero variance components: Shows the distribution of p-values from the *ad hoc* method described in the main text when variance components are zero under some representative scenarios.

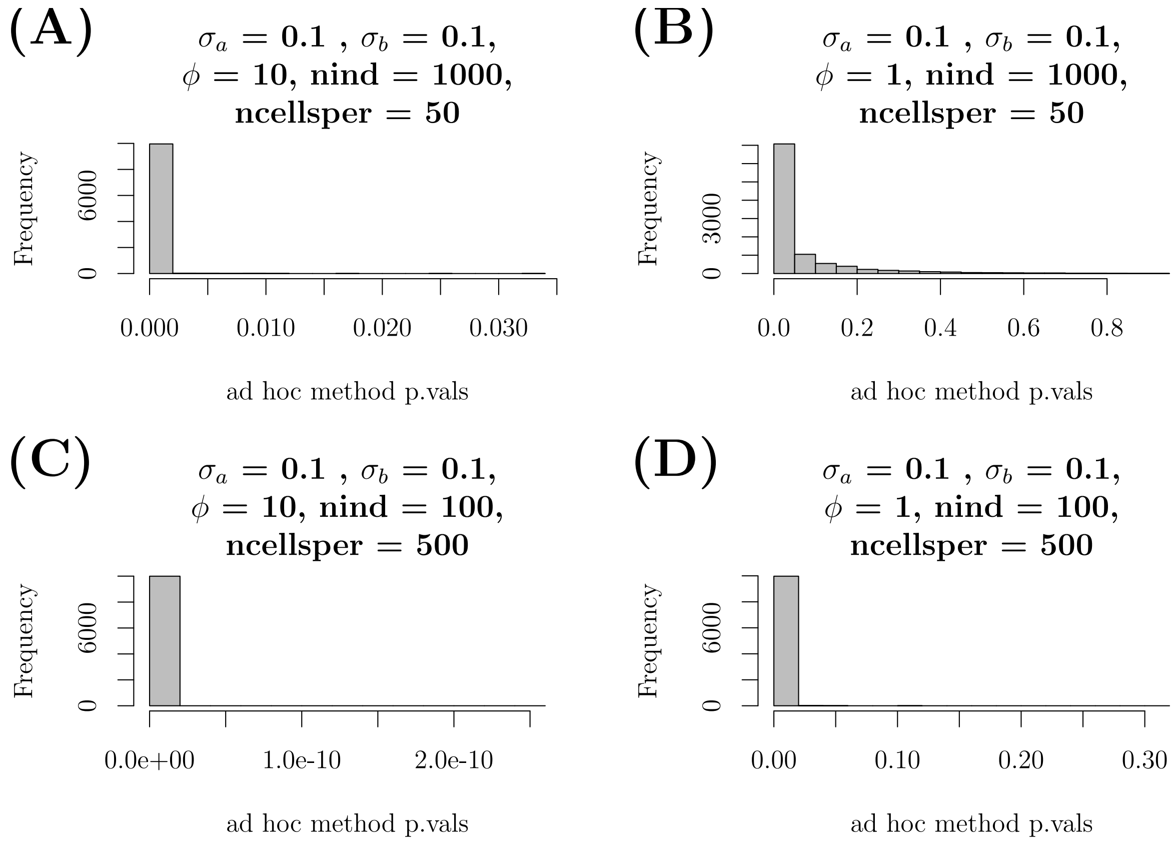


Supplementary Figure S3: *ad hoc* procedure with non-zero variance components: Shows the distribution of p-values from the *ad hoc* method described in section 3 of the main text when variance components are non-zero under some representative scenarios.

For each of the simulation results discussed in the main text and summarized in the tables below, we also calculated the p-values from the *ad hoc* method for determining if random effects are needed. Figures S2 and S3 show histograms of p-values from representative scenarios when variance components are zero and non-zero, respectively. When variance components are zero, p-values are close to uniformly distributed, meaning that most genes will not be flagged as in need of random effects. When variance components are non-zero, the method produces small p-values which can successfully flag the need to include random effects using a p-value cutoff threshold of, for example, 0.05 or 0.10.

### 3 More Detailed Type-I Error Simulation Results

Tables S1–S4 give more detailed results for the type-I error simulations at the 0.05 level. Each setting has results from three models: TWO-SIGMA, the zero-inflated negative binomial model (without random effects) "ZINB," and MAST [3]. Parameters $\alpha_1$ and $\beta_1$ correspond to the coefficients on a binary disease status indicator, and are set to 0 under the null and are non-zero under some alternative hypothesis. Other coefficients in both components include an intercept $(\alpha_0, \beta_0)$, coefficients from simulated age values $(\alpha_2, \beta_2)$ and coefficients from simulated CDR values $(\alpha_3, \beta_3)$. Parameter values were designed to mimic realistic values observed in the pancreas data analysis. "LRT" refers to the likelihood ratio statistic (on 2 d.f.), and the combined $\chi^2$ statistic is defined as the sum of the squared z-statistics from each of the two coefficients related to the binary disease status indicator. Coverage is given for parameters $\alpha_1$ and $\beta_1$, $\phi$, and $\sigma_a$ and $\sigma_b$. Note that confidence intervals for the variance components are computed on the log scale and exponentiated. Therefore, the intervals will not contain zero and thus coverage when $\sigma_a$ and/or $\sigma_b$ equal zero is not entirely meaningful. Finally, note that the average time column includes the average over all genes of the time needed for two runs of TWO-SIGMA, MAST, and the ZINB model (each with and without the coefficients corresponding to the binary disease status indicator) and to simulate the data as well as process the results for the entire replication. The purpose of these times is to highlight how total computation time varies within a table as variance component sizes change. Within a given table, differences in the running time are largely if not entirely due to TWO-SIGMA run time. More discussion of run time is given in the paragraph below.

For example, consider table S1. The first column "N/N Max" gives the number of genes that converged compared to the total number of genes simulated for each scenario, and as mentioned above the last column gives total runtime for all computations in a given simulation replication. One highly consistent trend is that the convergence percentage is lower and running time higher when variance components were zero. This is because the marginal likelihood for TWO-SIGMA is evaluated more times when the true values of $\sigma_a$ and $\sigma_b$ are on the boundary space of zero. Comparing the first and last runtimes in table S1 shows that there can be nearly a 50% increase in run-time when true variance components are zero yet random effects are included in the model. This underscores the usefulness of our *ad hoc* procedure to avoid fitting random effects where they are unnecessary. In table 1, Type-I error for TWO-SIGMA is well-preserved for any variance component value but becomes increasingly inflated for the ZINB model and MAST when variance components are non-zero. Furthermore, coverage from TWO-SIGMA for all parameters shown remains near the nominal level of 95%. These results hold well for all four cases varying the breakdown of the total of 50,000 cells between number of individuals and number of cells per individual, with a slight inflation of type-I error seen in table S4 and discussed in the main text.

Figures S4 and S5 show the observed type-I error across more stringent significance levels for representative simulation scenarios. These figures do not show systematically different patterns than seen at the .05 level in tables S1–S4 but can highlight the substantial type-I error seen for MAST and the ZINB model in some situations.

### Case 1: 1000 individuals, 50 single-cells each, 0.05 level

| N / N Max | Model | LRT Type-I Error | Combined $\chi^2$ Type-I Error | 95 % CI Coverage $\alpha_1$ | $\beta_1$ | $\phi$ | $\sigma_a$ | $\sigma_b$ | Simulation Parameters $\boldsymbol{\alpha} = (\alpha_0, \alpha_1, \alpha_2, \alpha_3)$ | $\boldsymbol{\beta} = (\beta_0, \beta_1, \beta_2, \beta_3)$ | $\phi$ | $\sigma_a$ | $\sigma_b$ | Avg. Time (min) |
|---|---|---|---|---|---|---|---|---|---|---|---|---|---|---|
| | TWO-SIGMA | **0.049** | **0.049** | 0.951 | 0.953 | 0.954 | — | — | | | | | | |
| 9195/10000 | ZINB | 0.051 | 0.051 | 0.950 | 0.951 | 0.953 | — | — | (1, **0**, -0.5, -2) | (2, **0**, -0.1, 0.6) | 10 | 0 | 0 | 31.7 |
| | MAST | 0.089 | 0.020 | 0.950 | 0.997 | — | — | — | | | | | | |
| | TWO-SIGMA | **0.048** | **0.047** | 0.953 | 0.951 | 0.952 | — | — | | | | | | |
| 9612/10000 | ZINB | 0.051 | 0.049 | 0.951 | 0.949 | 0.953 | — | — | (1, **0**, -0.5, -2) | (2, **0**, -0.1, 0.6) | 2 | 0 | 0 | 33.5 |
| | MAST | 0.080 | 0.032 | 0.950 | 0.978 | — | — | — | | | | | | |
| | TWO-SIGMA | **0.048** | **0.050** | 0.954 | 0.953 | 0.949 | — | — | | | | | | |
| 9464/10000 | ZINB | 0.052 | 0.053 | 0.952 | 0.951 | 0.949 | — | — | (1, **0**, -0.5, -2) | (2, **0**, -0.1, 0.6) | 1 | 0 | 0 | 32.2 |
| | MAST | 0.081 | 0.042 | 0.953 | 0.966 | — | — | — | | | | | | |
| | TWO-SIGMA | **0.051** | **0.051** | 0.949 | 0.952 | 0.952 | 0.936 | 0.948 | | | | | | |
| 10000/10000 | ZINB | 0.132 | 0.131 | 0.941 | 0.853 | 0.001 | — | — | (1, **0**, -0.5, -2) | (2, **0**, -0.1, 0.6) | 10 | 0.1 | 0.1 | 30.7 |
| | MAST | 0.144 | 0.059 | 0.942 | 0.950 | — | — | — | | | | | | |
| | TWO-SIGMA | **0.051** | **0.051** | 0.950 | 0.949 | 0.951 | 0.936 | 0.963 | | | | | | |
| 10000/10000 | ZINB | 0.078 | 0.078 | 0.945 | 0.918 | 0.666 | — | — | (1, **0**, -0.5, -2) | (2, **0**, -0.1, 0.6) | 2 | 0.1 | 0.1 | 30.7 |
| | MAST | 0.089 | 0.048 | 0.944 | 0.960 | — | — | — | | | | | | |
| | TWO-SIGMA | **0.049** | **0.049** | 0.948 | 0.948 | 0.950 | 0.935 | 0.954 | | | | | | |
| 10000/10000 | ZINB | 0.066 | 0.066 | 0.941 | 0.930 | 0.869 | — | — | (1, **0**, -0.5, -2) | (2, **0**, -0.1, 0.6) | 1 | 0.1 | 0.1 | 30.1 |
| | MAST | 0.095 | 0.049 | 0.941 | 0.960 | — | — | — | | | | | | |
| | TWO-SIGMA | **0.051** | **0.051** | 0.947 | 0.952 | 0.946 | 0.944 | 0.949 | | | | | | |
| 10000/10000 | ZINB | 0.621 | 0.621 | 0.776 | 0.417 | 0 | — | — | (1, **0**, -0.5, -2) | (2, **0**, -0.1, 0.6) | 10 | 0.5 | 0.5 | 20.9 |
| | MAST | 0.290 | 0.494 | 0.778 | 0.556 | — | — | — | | | | | | |
| | TWO-SIGMA | **0.053** | **0.053** | 0.948 | 0.948 | 0.947 | 0.947 | 0.946 | | | | | | |
| 9999/10000 | ZINB | 0.505 | 0.503 | 0.794 | 0.539 | 0 | — | — | (1, **0**, -0.5, -2) | (2, **0**, -0.1, 0.6) | 2 | 0.5 | 0.5 | 18.0 |
| | MAST | 0.275 | 0.400 | 0.792 | 0.658 | — | — | — | | | | | | |
| | TWO-SIGMA | **0.050** | **0.050** | 0.950 | 0.954 | 0.949 | 0.950 | 0.948 | | | | | | |
| 10000/10000 | ZINB | 0.404 | 0.398 | 0.817 | 0.639 | 0 | — | — | (1, **0**, -0.5, -2) | (2, **0**, -0.1, 0.6) | 1 | 0.5 | 0.5 | 17.0 |
| | MAST | 0.247 | 0.301 | 0.810 | 0.759 | — | — | — | | | | | | |

Supplementary Table S1: Type-I Error using LRT to test $H_0 : \alpha_1 = 0,\ \beta_1 = 0$ with a significance level of 0.05

### Case 2: 500 individuals, 100 single-cells each, 0.05 level

| N / N Max | Model | LRT Type-I Error | Combined $\chi^2$ Type-I Error | 95 % CI Coverage $\alpha_1$ | $\beta_1$ | $\phi$ | $\sigma_a$ | $\sigma_b$ | Simulation Parameters $\boldsymbol{\alpha} = (\alpha_0, \alpha_1, \alpha_2, \alpha_3)$ | $\boldsymbol{\beta} = (\beta_0, \beta_1, \beta_2, \beta_3)$ | $\phi$ | $\sigma_a$ | $\sigma_b$ | Avg. Time (min) |
|---|---|---|---|---|---|---|---|---|---|---|---|---|---|---|
| | TWO-SIGMA | **0.044** | **0.044** | 0.955 | 0.952 | 0.950 | — | — | | | | | | |
| 8895/10000 | ZINB | 0.047 | 0.047 | 0.953 | 0.950 | 0.952 | — | — | (1, **0**, -0.5, -2) | (2, **0**, -0.1, 0.6) | 10 | 0 | 0 | 31.8 |
| | MAST | 0.086 | 0.020 | 0.953 | 0.996 | — | — | — | | | | | | |
| | TWO-SIGMA | **0.042** | **0.043** | 0.952 | 0.957 | 0.946 | — | — | | | | | | |
| 9285/10000 | ZINB | 0.045 | 0.046 | 0.949 | 0.955 | 0.949 | — | — | (1, **0**, -0.5, -2) | (2, **0**, -0.1, 0.6) | 2 | 0 | 0 | 32.7 |
| | MAST | 0.084 | 0.030 | 0.949 | 0.981 | — | — | — | | | | | | |
| | TWO-SIGMA | **0.046** | **0.045** | 0.952 | 0.952 | 0.952 | — | — | | | | | | |
| 9502/10000 | ZINB | 0.049 | 0.049 | 0.951 | 0.949 | 0.951 | — | — | (1, **0**, -0.5, -2) | (2, **0**, -0.1, 0.6) | 1 | 0 | 0 | 30.6 |
| | MAST | 0.085 | 0.039 | 0.950 | 0.969 | — | — | — | | | | | | |
| | TWO-SIGMA | **0.052** | **0.052** | 0.942 | 0.951 | 0.950 | 0.955 | 0.948 | | | | | | |
| 10000/10000 | ZINB | 0.217 | 0.218 | 0.926 | 0.754 | 0.002 | — | — | (1, **0**, -0.5, -2) | (2, **0**, -0.1, 0.6) | 10 | 0.1 | 0.1 | 23.8 |
| | MAST | 0.187 | 0.099 | 0.926 | 0.899 | — | — | — | | | | | | |
| | TWO-SIGMA | **0.050** | **0.052** | 0.948 | 0.953 | 0.950 | 0.948 | 0.958 | | | | | | |
| 10000/10000 | ZINB | 0.106 | 0.105 | 0.935 | 0.885 | 0.670 | — | — | (1, **0**, -0.5, -2) | (2, **0**, -0.1, 0.6) | 2 | 0.1 | 0.1 | 25.3 |
| | MAST | 0.104 | 0.065 | 0.934 | 0.944 | — | — | — | | | | | | |
| | TWO-SIGMA | **0.052** | **0.052** | 0.950 | 0.950 | 0.949 | 0.946 | 0.967 | | | | | | |
| 10000/10000 | ZINB | 0.081 | 0.082 | 0.939 | 0.914 | 0.866 | — | — | (1, **0**, -0.5, -2) | (2, **0**, -0.1, 0.6) | 1 | 0.1 | 0.1 | 22.9 |
| | MAST | 0.107 | 0.063 | 0.938 | 0.945 | — | — | — | | | | | | |
| | TWO-SIGMA | **0.051** | **0.052** | 0.950 | 0.949 | 0.946 | 0.948 | 0.947 | | | | | | |
| 10000/10000 | ZINB | 0.753 | 0.753 | 0.667 | 0.307 | 0 | — | — | (1, **0**, -0.5, -2) | (2, **0**, -0.1, 0.6) | 10 | 0.5 | 0.5 | 17.8 |
| | MAST | 0.419 | 0.673 | 0.670 | 0.414 | — | — | — | | | | | | |
| | TWO-SIGMA | **0.054** | **0.055** | 0.951 | 0.942 | 0.953 | 0.949 | 0.950 | | | | | | |
| 10000/10000 | ZINB | 0.663 | 0.663 | 0.691 | 0.414 | 0 | — | — | (1, **0**, -0.5, -2) | (2, **0**, -0.1, 0.6) | 2 | 0.5 | 0.5 | 18.0 |
| | MAST | 0.382 | 0.579 | 0.685 | 0.516 | — | — | — | | | | | | |
| | TWO-SIGMA | **0.054** | **0.054** | 0.944 | 0.948 | 0.947 | 0.950 | 0.942 | | | | | | |
| 10000/10000 | ZINB | 0.583 | 0.577 | 0.705 | 0.509 | 0 | — | — | (1, **0**, -0.5, -2) | (2, **0**, -0.1, 0.6) | 1 | 0.5 | 0.5 | 15.5 |
| | MAST | 0.366 | 0.494 | 0.696 | 0.628 | — | — | — | | | | | | |

Supplementary Table S2: Type-I Error using LRT to test $H_0 : \alpha_1 = 0,\ \beta_1 = 0$ with a significance level of 0.05

### Case 3: 100 individuals, 500 single-cells each, 0.05 level

| | | LRT | Combined $\chi^2$ | 95 % CI Coverage | | | | | Simulation Parameters | | | | | Avg. Time |
|---|---|---|---|---|---|---|---|---|---|---|---|---|---|---|
| N / N Max | Model | Type-I Error | Type-I Error | $\alpha_1$ | $\beta_1$ | $\phi$ | $\sigma_a$ | $\sigma_b$ | $\boldsymbol{\alpha} = (\alpha_0, \alpha_1, \alpha_2, \alpha_3)$ | $\boldsymbol{\beta} = (\beta_0, \beta_1, \beta_2, \beta_3)$ | $\phi$ | $\sigma_a$ | $\sigma_b$ | (min) |
| | TWO-SIGMA | **0.042** | **0.044** | 0.954 | 0.953 | 0.951 | — | — | | | | | | |
| 8773/10000 | ZINB | 0.050 | 0.050 | 0.950 | 0.950 | 0.950 | — | — | (1, **0**, -0.5, -2) | (2, **0**, -0.1, 0.6) | 10 | 0 | 0 | 32.1 |
| | MAST | 0.090 | 0.021 | 0.950 | 0.995 | — | — | — | | | | | | |
| | TWO-SIGMA | **0.038** | **0.038** | 0.960 | 0.957 | 0.953 | — | — | | | | | | |
| 8901/10000 | ZINB | 0.044 | 0.043 | 0.955 | 0.954 | 0.953 | — | — | (1, **0**, -0.5, -2) | (2, **0**, -0.1, 0.6) | 2 | 0 | 0 | 32.1 |
| | MAST | 0.079 | 0.028 | 0.955 | 0.977 | — | — | — | | | | | | |
| | TWO-SIGMA | **0.044** | **0.045** | 0.956 | 0.954 | 0.951 | — | — | | | | | | |
| 9199/10000 | ZINB | 0.051 | 0.050 | 0.952 | 0.949 | 0.951 | — | — | (1, **0**, -0.5, -2) | (2, **0**, -0.1, 0.6) | 1 | 0 | 0 | 31.3 |
| | MAST | 0.087 | 0.038 | 0.950 | 0.969 | — | — | — | | | | | | |
| | TWO-SIGMA | **0.056** | **0.059** | 0.938 | 0.947 | 0.951 | 0.979 | 0.938 | | | | | | |
| 9999/10000 | ZINB | 0.534 | 0.533 | 0.869 | 0.465 | 0.007 | — | — | (1, **0**, -0.5, -2) | (2, **0**, -0.1, 0.6) | 10 | 0.1 | 0.1 | 25.7 |
| | MAST | 0.313 | 0.376 | 0.869 | 0.634 | — | — | — | | | | | | |
| | TWO-SIGMA | **0.057** | **0.060** | 0.940 | 0.942 | 0.952 | 0.978 | 0.943 | | | | | | |
| 10000/10000 | ZINB | 0.323 | 0.320 | 0.877 | 0.685 | 0.673 | — | — | (1, **0**, -0.5, -2) | (2, **0**, -0.1, 0.6) | 2 | 0.1 | 0.1 | 25.9 |
| | MAST | 0.176 | 0.226 | 0.872 | 0.791 | — | — | — | | | | | | |
| | TWO-SIGMA | **0.053** | **0.058** | 0.939 | 0.947 | 0.950 | 0.977 | 0.955 | | | | | | |
| 10000/10000 | ZINB | 0.224 | 0.219 | 0.887 | 0.789 | 0.883 | — | — | (1, **0**, -0.5, -2) | (2, **0**, -0.1, 0.6) | 1 | 0.1 | 0.1 | 25.6 |
| | MAST | 0.174 | 0.169 | 0.882 | 0.860 | — | — | — | | | | | | |
| | TWO-SIGMA | **0.055** | **0.058** | 0.945 | 0.942 | 0.951 | 0.935 | 0.936 | | | | | | |
| 10000/10000 | ZINB | 0.941 | 0.941 | 0.367 | 0.142 | 0 | — | — | (1, **0**, -0.5, -2) | (2, **0**, -0.1, 0.6) | 10 | 0.5 | 0.5 | 21.4 |
| | MAST | 0.716 | 0.914 | 0.367 | 0.193 | — | — | — | | | | | | |
| | TWO-SIGMA | **0.056** | **0.060** | 0.940 | 0.945 | 0.950 | 0.936 | 0.934 | | | | | | |
| 10000/10000 | ZINB | 0.909 | 0.909 | 0.386 | 0.196 | 0 | — | — | (1, **0**, -0.5, -2) | (2, **0**, -0.1, 0.6) | 2 | 0.5 | 0.5 | 20.4 |
| | MAST | 0.685 | 0.884 | 0.383 | 0.254 | — | — | — | | | | | | |
| | TWO-SIGMA | **0.053** | **0.056** | 0.943 | 0.947 | 0.952 | 0.939 | 0.934 | | | | | | |
| 10000/10000 | ZINB | 0.873 | 0.872 | 0.412 | 0.256 | 0 | — | — | (1, **0**, -0.5, -2) | (2, **0**, -0.1, 0.6) | 1 | 0.5 | 0.5 | 20.0 |
| | MAST | 0.649 | 0.839 | 0.400 | 0.324 | — | — | — | | | | | | |

Supplementary Table S3: Type-I Error using LRT to test $H_0 : \alpha_1 = 0,\ \beta_1 = 0$ with a significance level of 0.05

### Case 4: 25 individuals, 2000 single-cells each, 0.05 level

| | | LRT | Combined $\chi^2$ | 95 % CI Coverage | | | | | Simulation Parameters | | | | | Avg. Time |
|---|---|---|---|---|---|---|---|---|---|---|---|---|---|---|
| N / N Max | Model | Type-I Error | Type-I Error | $\alpha_1$ | $\beta_1$ | $\phi$ | $\sigma_a$ | $\sigma_b$ | $\boldsymbol{\alpha} = (\alpha_0, \alpha_1, \alpha_2, \alpha_3)$ | $\boldsymbol{\beta} = (\beta_0, \beta_1, \beta_2, \beta_3)$ | $\phi$ | $\sigma_a$ | $\sigma_b$ | (min) |
| | TWO-SIGMA | **0.041** | **0.045** | 0.953 | 0.951 | 0.950 | — | — | | | | | | |
| 8698/10000 | ZINB | 0.052 | 0.052 | 0.947 | 0.946 | 0.950 | — | — | (1, **0**, -0.5, -2) | (2, **0**, -0.1, 0.6) | 10 | 0 | 0 | 28.9 |
| | MAST | 0.090 | 0.021 | 0.947 | 0.995 | — | — | — | | | | | | |
| | TWO-SIGMA | **0.041** | **0.046** | 0.955 | 0.954 | 0.952 | — | — | | | | | | |
| 8585/10000 | ZINB | 0.052 | 0.053 | 0.949 | 0.949 | 0.952 | — | — | (1, **0**, -0.5, -2) | (2, **0**, -0.1, 0.6) | 2 | 0 | 0 | 28.3 |
| | MAST | 0.086 | 0.034 | 0.948 | 0.976 | — | — | — | | | | | | |
| | TWO-SIGMA | **0.042** | **0.044** | 0.954 | 0.954 | 0.946 | — | — | | | | | | |
| 8763/10000 | ZINB | 0.051 | 0.050 | 0.949 | 0.949 | 0.946 | — | — | (1, **0**, -0.5, -2) | (2, **0**, -0.1, 0.6) | 1 | 0 | 0 | 27.8 |
| | MAST | 0.090 | 0.041 | 0.949 | 0.966 | — | — | — | | | | | | |
| | TWO-SIGMA | **0.076** | **0.088** | 0.920 | 0.923 | 0.949 | 0.980 | 0.909 | | | | | | |
| 9544/10000 | ZINB | 0.817 | 0.817 | 0.689 | 0.235 | 0.056 | — | — | (1, **0**, -0.5, -2) | (2, **0**, -0.1, 0.6) | 10 | 0.1 | 0.1 | 22.3 |
| | MAST | 0.497 | 0.720 | 0.689 | 0.354 | — | — | — | | | | | | |
| | TWO-SIGMA | **0.072** | **0.087** | 0.926 | 0.923 | 0.946 | 0.994 | 0.896 | | | | | | |
| 9999/10000 | ZINB | 0.643 | 0.642 | 0.708 | 0.424 | 0.719 | — | — | (1, **0**, -0.5, -2) | (2, **0**, -0.1, 0.6) | 2 | 0.1 | 0.1 | 26.4 |
| | MAST | 0.361 | 0.562 | 0.704 | 0.527 | — | — | — | | | | | | |
| | TWO-SIGMA | **0.075** | **0.094** | 0.922 | 0.923 | 0.949 | 0.992 | 0.906 | | | | | | |
| 10000/10000 | ZINB | 0.548 | 0.542 | 0.733 | 0.541 | 0.880 | — | — | (1, **0**, -0.5, -2) | (2, **0**, -0.1, 0.6) | 1 | 0.1 | 0.1 | 26.3 |
| | MAST | 0.361 | 0.467 | 0.718 | 0.637 | — | — | — | | | | | | |
| | TWO-SIGMA | **0.076** | **0.094** | 0.920 | 0.920 | 0.951 | 0.888 | 0.888 | | | | | | |
| 10000/10000 | ZINB | 0.984 | 0.984 | 0.194 | 0.070 | 0 | — | — | (1, **0**, -0.5, -2) | (2, **0**, -0.1, 0.6) | 10 | 0.5 | 0.5 | 22.7 |
| | MAST | 0.875 | 0.979 | 0.195 | 0.098 | — | — | — | | | | | | |
| | TWO-SIGMA | **0.076** | **0.092** | 0.925 | 0.922 | 0.949 | 0.886 | 0.882 | | | | | | |
| 10000/10000 | ZINB | 0.974 | 0.975 | 0.202 | 0.101 | 0 | — | — | (1, **0**, -0.5, -2) | (2, **0**, -0.1, 0.6) | 2 | 0.5 | 0.5 | 22.0 |
| | MAST | 0.857 | 0.966 | 0.197 | 0.132 | — | — | — | | | | | | |
| | TWO-SIGMA | **0.074** | **0.089** | 0.923 | 0.922 | 0.950 | 0.891 | 0.880 | | | | | | |
| 10000/10000 | ZINB | 0.964 | 0.963 | 0.218 | 0.135 | 0 | — | — | (1, **0**, -0.5, -2) | (2, **0**, -0.1, 0.6) | 1 | 0.5 | 0.5 | 22.5 |
| | MAST | 0.827 | 0.953 | 0.213 | 0.174 | — | — | — | | | | | | |

Supplementary Table S4: Type-I Error using LRT to test $H_0 : \alpha_1 = 0,\ \beta_1 = 0$ with a significance level of 0.05

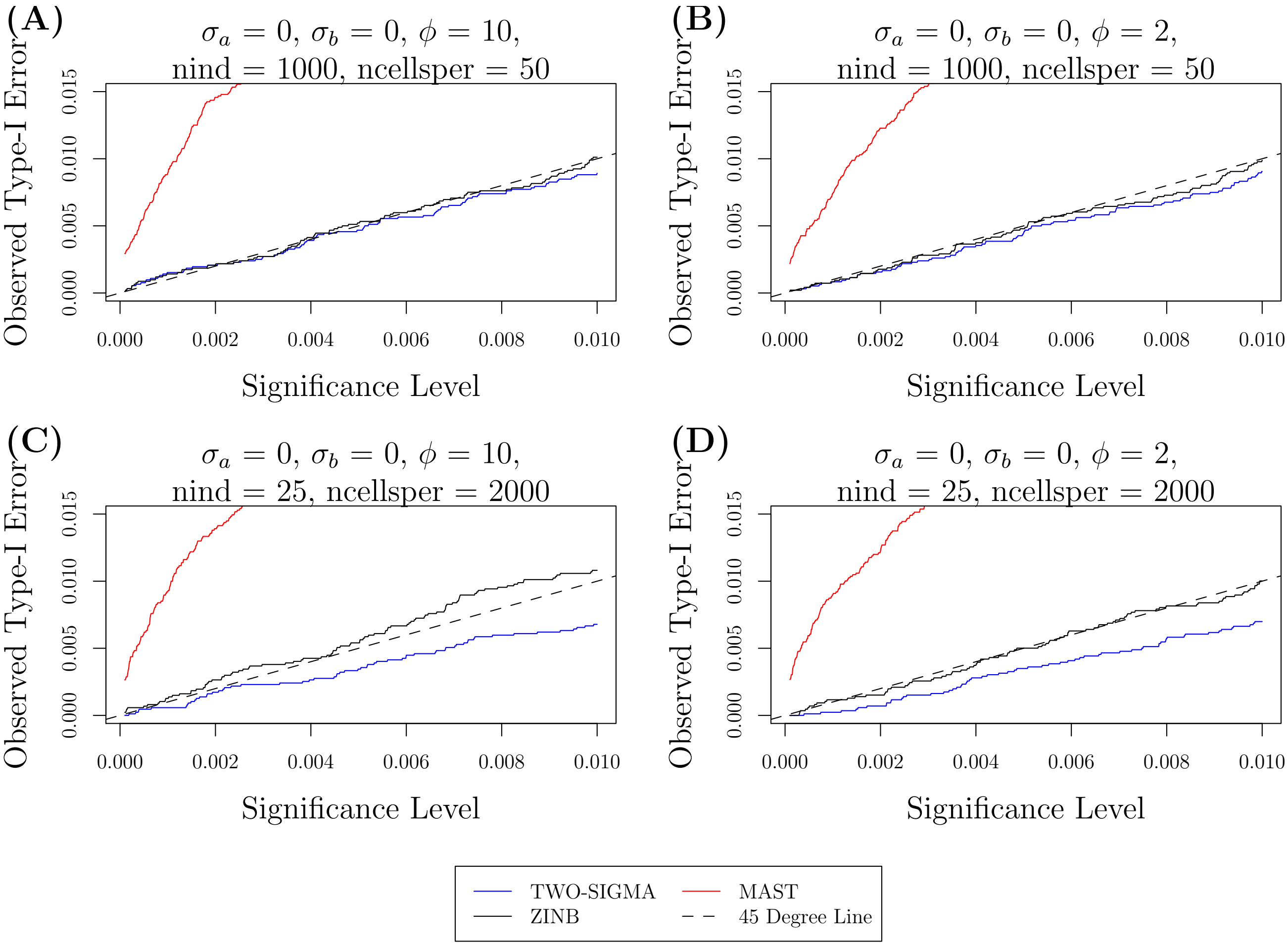


Supplementary Figure S4: Type-I error across different significance levels: Shows the observed type-I error across various nominal significance levels.

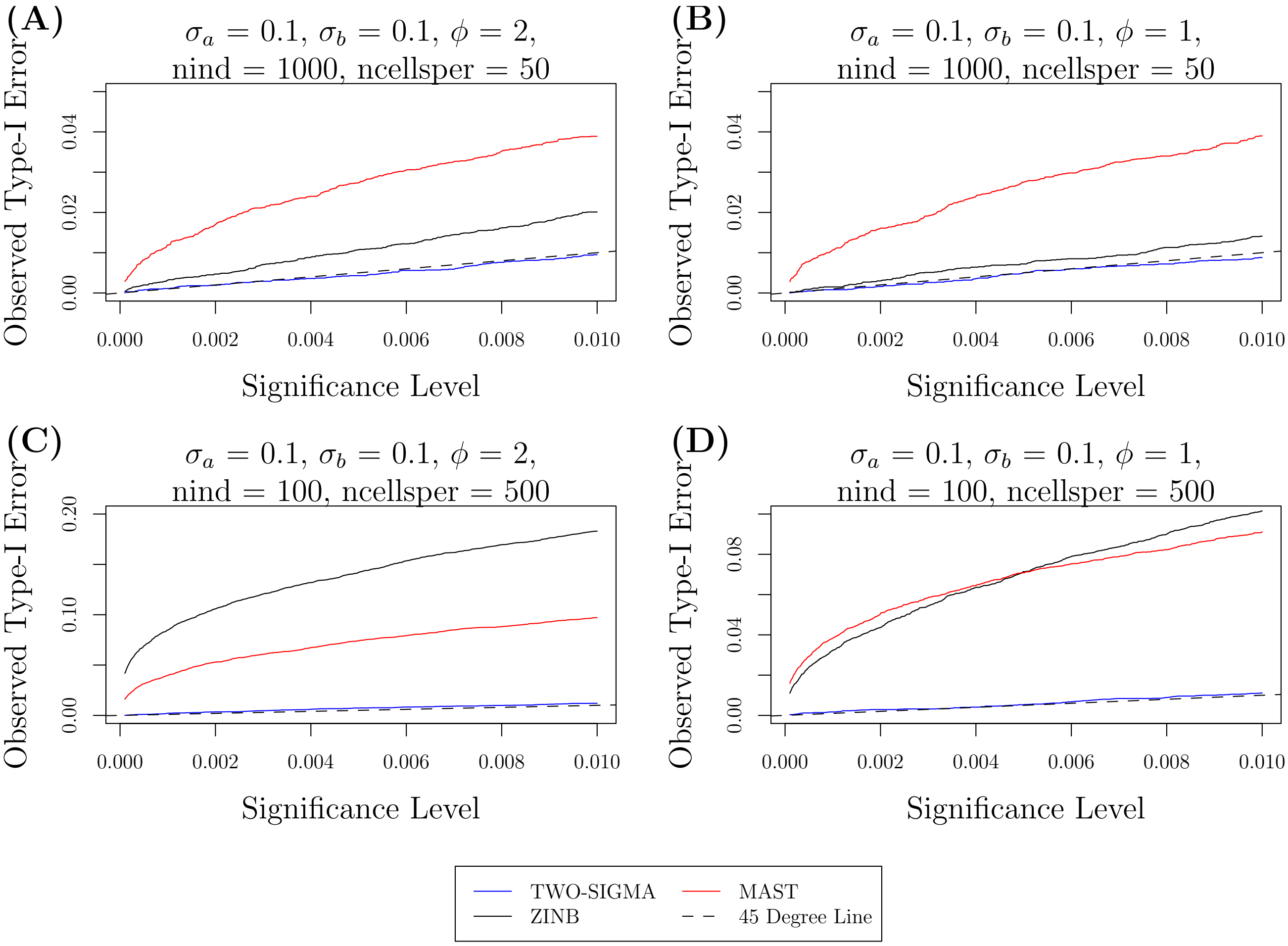


Supplementary Figure S5: Type-I error across different significance levels: Shows the observed type-I error across various nominal significance levels.

### 4 Power Results

Simulations under the same framework were also performed for non-zero values of $\alpha_1$ and $\beta_1$ (both as defined in the previous section) to evaluate the power of TWO-SIGMA in testing $\mathrm{H}_0 : \alpha_1 = 0$, $\beta_1 = 0$. As seen in Table 1 of the main text and tables S1–S4 of this supplement, MAST and the ZINB model can suffer from vastly inflated type-I error. Thus, the observed (or "apparent") power does not always provide a fair comparison to TWO-SIGMA. For each of the three methods we therefore calculated empirical significance thresholds for all null simulation settings. These are cutoffs such that the percentage of statistics larger than the threshold equals the significance level. "True" power is then calculated by rejecting the null if the test statistic is larger than the empirical significance threshold from the corresponding simulation setting under the null. In simulation settings this does not add computation, but in real data setting this procedure involves additional computation and is therefore not preferred.

Because the type-I error for TWO-SIGMA is approximately preserved in all four sample size cases, true power is nearly identical to apparent power for TWO-SIGMA. We therefore found it unnecessary to use true power for TWO-SIGMA in supplementary figures S6-S8 shown here and figure 2 in the main text. In contrast, true power can be very different than apparent power for both the ZINB model and MAST given their inflated type-I errors. For example, one simulation setting shows that the apparent power of MAST is 0.375 which the true power for this scenario is only 0.194 (see the third rows of table S5 and S9). Although not presented, this discrepancy between apparent and true power would be even more pronounced if the simulated data here were based on larger values of the variance components $\sigma_a$ and $\sigma_b$ because type-I errors are more inflated for larger variance components (see tables S1-S4).

One general observation from tables S9 to S12 and figure 2 of the main text is that the ZINB model retains very high true power in both sample size settings and across all four effect scenarios. For smaller values of $\alpha_0$ the ZINB model can sometimes have higher true power than TWO-SIGMA. As the dropout proportion increases (via increasing $\alpha_0$), TWO-SIGMA tends to eventually have higher power. TWO-SIGMA does not require the use of computationally expensive resampling procedures for valid inference, giving it a key advantage over the ZINB model, which is not articulated explicitly as a DE method for scRNA-seq data.

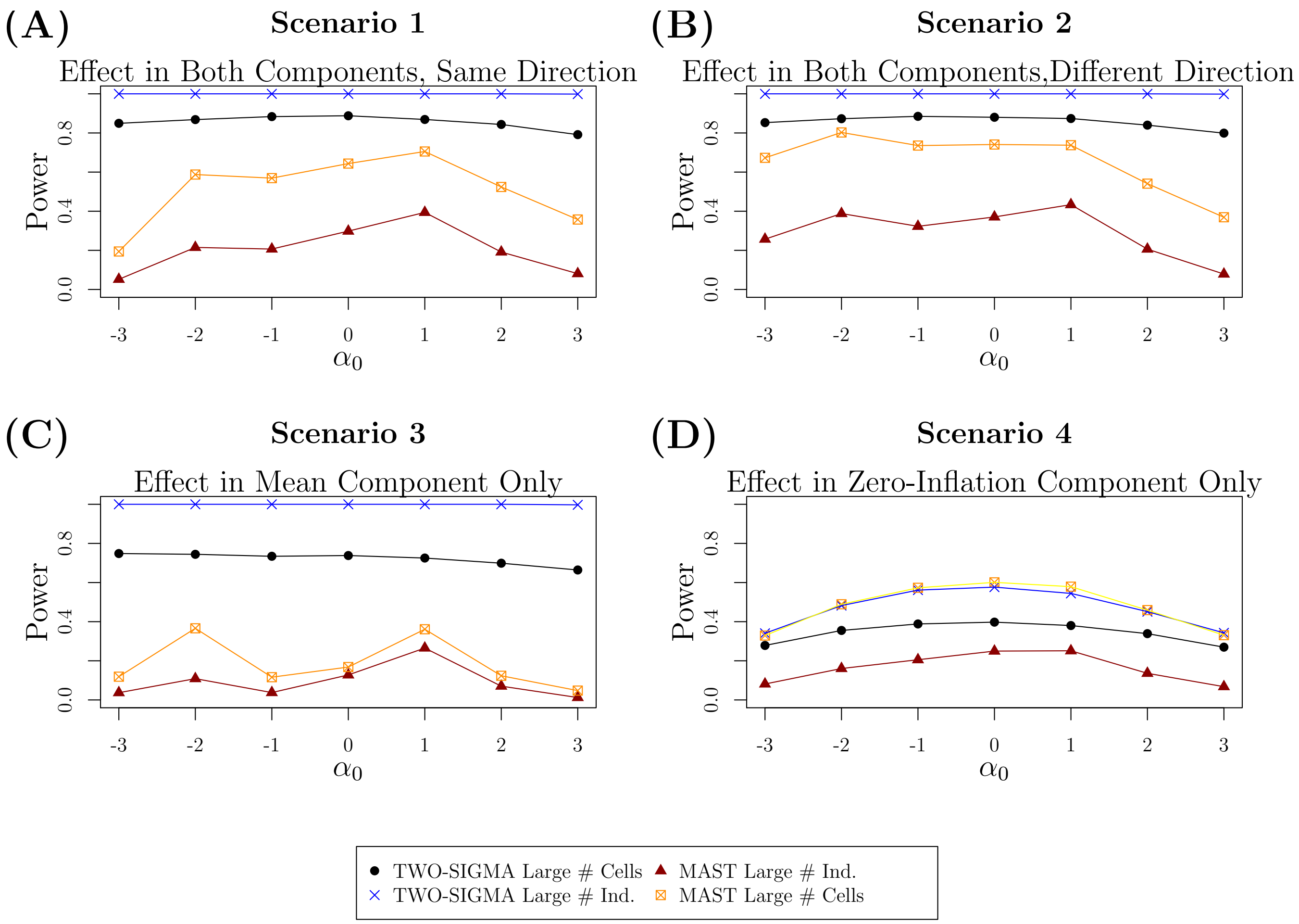


Supplementary Figure S6: Power evaluations in simulated data: Shows the power to test $H_0 : \alpha_1 = \beta_1 = 0$ by varying the intercept $\alpha_0$ to control the drop-out proportion in four setups: TWO-SIGMA and MAST with 50 cells from each of 1000 individuals or 500 cells from each of 100 individuals. Values of $\phi$, $\sigma_a$, and $\sigma_b$ were all set to 0.1 and an effect size of 0.03 was used. Larger values of $\alpha_0$ correspond to more drop-out in the data. 10,000 genes were simulated. Because of the type-I error inflation from MAST seen in tables S1–S4, true power was calculated and plotted using the empirical significance threshold from the corresponding setting under the null. TWO-SIGMA retains higher power in the first three scenarios and half of the fourth scenario without the need to use true power. See section 4 of the supplement for more details about computing true power and discussion regarding power trends across all three methods.

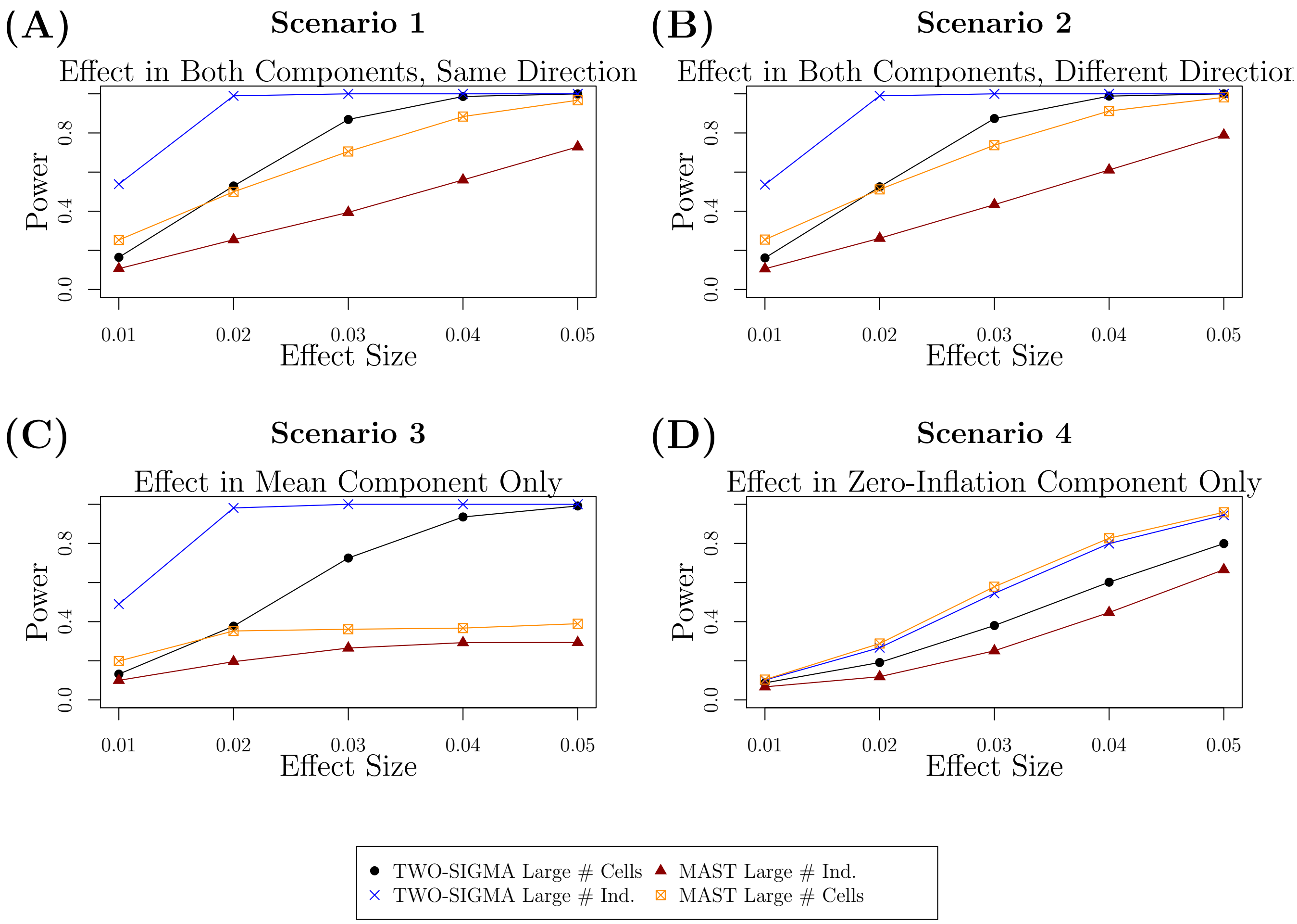


Supplementary Figure S7: Power evaluations in simulated data: Shows the power to test $H_0 : \alpha_1 = \beta_1 = 0$ by varying the effect size in two sample size setups: 50 cells from each of 1000 individuals or 500 cells from each of 100 individuals. Values of $\phi$, $\sigma_a$, and $\sigma_b$ were all set to 0.1 and 10,000 genes were simulated. Because of the type-I error inflation from the ZINB model and MAST seen in tables S1–S4, true power was calculated and plotted using the empirical significance threshold from the corresponding setting under the null for both of these methods. TWO-SIGMA retains higher power in the first three scenarios and half of the fourth scenario without the need to use true power. See the discussion at the beginning of section 4 of the supplement for more details about computing true power and discussion regarding power trends across all differing methods.

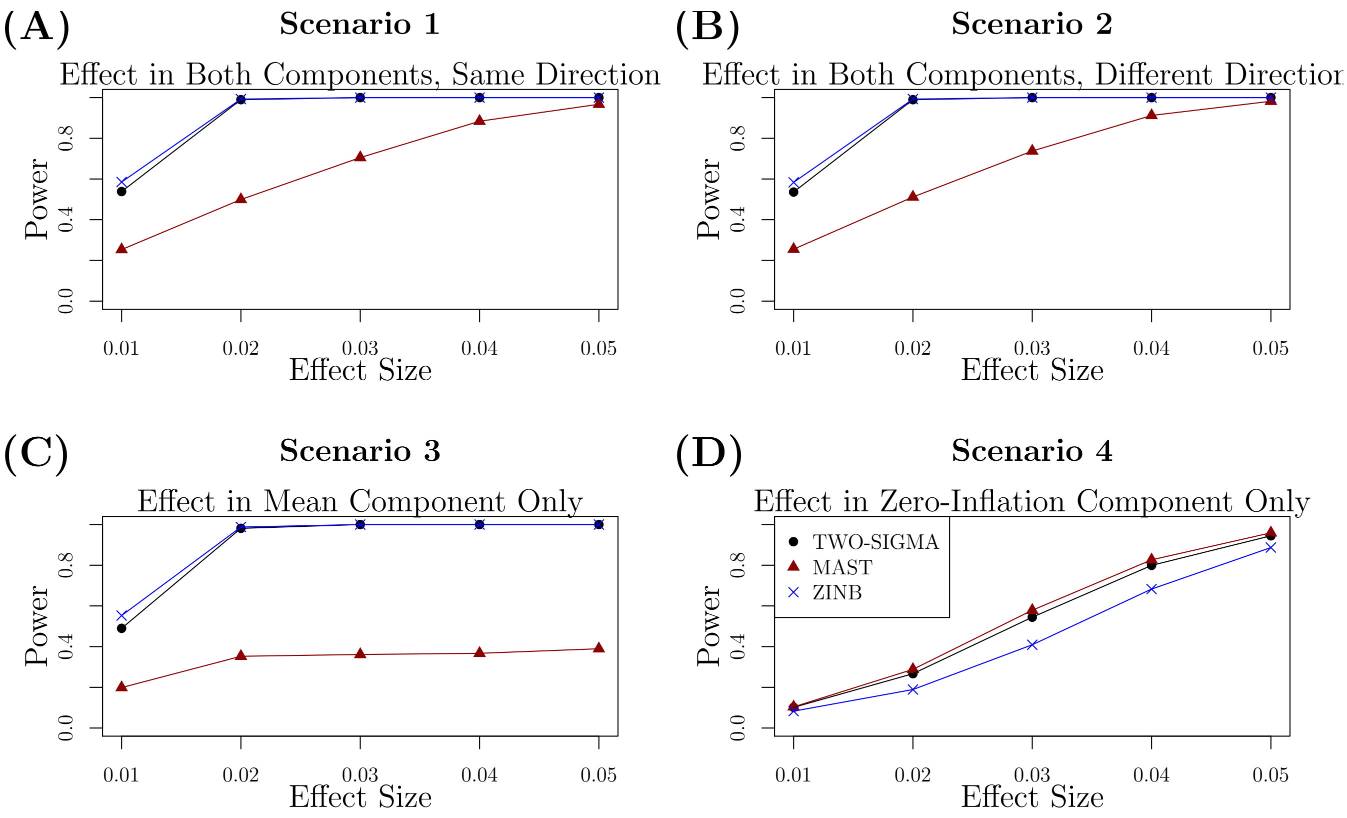


Supplementary Figure S8: Power evaluations in simulated data: Shows the power to test $H_0 : \alpha_1 = \beta_1 = 0$ by varying the effect size with 50 cells from each of 1000 individuals. Values of $\phi$, $\sigma_a$, and $\sigma_b$ were all set to 0.1 and 10,000 genes were simulated. Because of the type-I error inflation from the ZINB model and MAST seen in tables S1–S4, true power was calculated and plotted using the empirical significance threshold from the corresponding setting under the null for these two methods. In the first three scenarios, MAST consistently has lower true power while TWO-SIGMA and the ZINB model typically have very similar true power. When the effect is only in the zero-inflation component, power is lower for all methods at all effect sizes. Using TWO-SIGMA can bypass the need for computationally expensive resampling procedures needed to generate true power. See the discussion at the beginning of section 4 of the supplement for more details about computing true power and discussion regarding power trends across all differing methods.

#### 4.1 Results using "Apparent" Power for MAST and the ZINB model

### Case 1: 1000 individuals, 50 single-cells each, 0.05 level

##### Power Scenarios 1 & 2 from Figure 2 of main text

##### Effects in both components in either the same or different directions

| | | LRT | Combined $\chi^2$ | Simulation Parameters | | | | | Avg. Time |
|---|---|---|---|---|---|---|---|---|---|
| N / N Max | Model | P(Reject $H_0$) | P(Reject $H_0$) | $\boldsymbol{\alpha} = (\alpha_0, \alpha_1, \alpha_2, \alpha_3)$ | $\boldsymbol{\beta} = (\beta_0, \beta_1, \beta_2, \beta_3)$ | $\phi$ | $\sigma_a$ | $\sigma_b$ | (min) |
| | TWO-SIGMA | **1.000** | **1.000** | | | | | | |
| 10000/10000 | ZINB | 1.000 | 1.000 | (-3, **0.03**, -0.5, -2) | (2, **0.03**, -0.1, 0.6) | 10 | 0.1 | 0.1 | 27.9 |
| | MAST | 0.375 | 1.000 | | | | | | |
| | TWO-SIGMA | **1.000** | **1.000** | | | | | | |
| 10000/10000 | ZINB | 1.000 | 1.000 | (-2, **0.03**, -0.5, -2) | (2, **0.03**, -0.1, 0.6) | 10 | 0.1 | 0.1 | 27.8 |
| | MAST | 0.738 | 1.000 | | | | | | |
| | TWO-SIGMA | **1.000** | **1.000** | | | | | | |
| 10000/10000 | ZINB | 1.000 | 1.000 | (-1, **0.03**, -0.5, -2) | (2, **0.03**, -0.1, 0.6) | 10 | 0.1 | 0.1 | 27.0 |
| | MAST | 0.767 | 1.000 | | | | | | |
| | TWO-SIGMA | **1.000** | **1.000** | | | | | | |
| 10000/10000 | ZINB | 1.000 | 1.000 | (0, **0.03**, -0.5, -2) | (2, **0.03**, -0.1, 0.6) | 10 | 0.1 | 0.1 | 29.4 |
| | MAST | 0.816 | 1.000 | | | | | | |
| | TWO-SIGMA | **1.000** | **1.000** | | | | | | |
| 10000/10000 | ZINB | 1.000 | 1.000 | (1, **0.03**, -0.5, -2) | (2, **0.03**, -0.1, 0.6) | 10 | 0.1 | 0.1 | 29.4 |
| | MAST | 0.834 | 1.000 | | | | | | |
| | TWO-SIGMA | **1.000** | **1.000** | | | | | | |
| 10000/10000 | ZINB | 1.000 | 1.000 | (2, **0.03**, -0.5, -2) | (2, **0.03**, -0.1, 0.6) | 10 | 0.1 | 0.1 | 27.5 |
| | MAST | 0.720 | 0.999 | | | | | | |
| | TWO-SIGMA | **0.999** | **0.999** | | | | | | |
| 10000/10000 | ZINB | 1.000 | 1.000 | (3, **0.03**, -0.5, -2) | (2, **0.03**, -0.1, 0.6) | 10 | 0.1 | 0.1 | 27.2 |
| | MAST | 0.572 | 0.986 | | | | | | |
| | TWO-SIGMA | **1.000** | **1.000** | | | | | | |
| 10000/10000 | ZINB | 1.000 | 1.000 | (-3, **0.03**, -0.5, -2) | (2, **-0.03**, -0.1, 0.6) | 10 | 0.1 | 0.1 | 27.4 |
| | MAST | 0.843 | 1.000 | | | | | | |
| | TWO-SIGMA | **1.000** | **1.000** | | | | | | |
| 10000/10000 | ZINB | 1.000 | 1.000 | (-2, **0.03**, -0.5, -2) | (2, **-0.03**, -0.1, 0.6) | 10 | 0.1 | 0.1 | 27.6 |
| | MAST | 0.913 | 1.000 | | | | | | |
| | TWO-SIGMA | **1.000** | **1.000** | | | | | | |
| 10000/10000 | ZINB | 1.000 | 1.000 | (-1, **0.03**, -0.5, -2) | (2, **-0.03**, -0.1, 0.6) | 10 | 0.1 | 0.1 | 27.7 |
| | MAST | 0.884 | 1.000 | | | | | | |
| | TWO-SIGMA | **1.000** | **1.000** | | | | | | |
| 10000/10000 | ZINB | 1.000 | 1.000 | (0, **0.03**, -0.5, -2) | (2, **-0.03**, -0.1, 0.6) | 10 | 0.1 | 0.1 | 29.5 |
| | MAST | 0.879 | 1.000 | | | | | | |
| | TWO-SIGMA | **1.000** | **1.000** | | | | | | |
| 10000/10000 | ZINB | 1.000 | 1.000 | (1, **0.03**, -0.5, -2) | (2, **-0.03**, -0.1, 0.6) | 10 | 0.1 | 0.1 | 28.2 |
| | MAST | 0.863 | 1.000 | | | | | | |
| | TWO-SIGMA | **1.000** | **1.000** | | | | | | |
| 10000/10000 | ZINB | 1.000 | 1.000 | (2, **0.03**, -0.5, -2) | (2, **-0.03**, -0.1, 0.6) | 10 | 0.1 | 0.1 | 27.0 |
| | MAST | 0.734 | 0.999 | | | | | | |
| | TWO-SIGMA | **0.999** | **0.999** | | | | | | |
| 10000/10000 | ZINB | 0.999 | 0.999 | (3, **0.03**, -0.5, -2) | (2, **-0.03**, -0.1, 0.6) | 10 | 0.1 | 0.1 | 26.2 |
| | MAST | 0.579 | 0.985 | | | | | | |

Supplementary Table S5: Apparent Power using LRT to test $H_0 : \alpha_1 = 0, \beta_1 = 0$ with a significance level of 0.05

### Case 1: 1000 individuals, 50 single-cells each, 0.05 level

##### Power Scenarios 3 & 4 from Figure 2 of main text

##### Effects in one component at a time

| N / N Max | Model | LRT P(Reject $H_0$) | Combined $\chi^2$ P(Reject $H_0$) | Simulation Parameters $\boldsymbol{\alpha} = (\alpha_0, \alpha_1, \alpha_2, \alpha_3)$ | $\boldsymbol{\beta} = (\beta_0, \beta_1, \beta_2, \beta_3)$ | $\phi$ | $\sigma_a$ | $\sigma_b$ | Avg. Time (min) |
|---|---|---|---|---|---|---|---|---|---|
| | TWO-SIGMA | **1.000** | **1.000** | | | | | | |
| 10000/10000 | ZINB | 1.000 | 1.000 | (-3, **0**, -0.5, -2) | (2, **0.03**, -0.1, 0.6) | 10 | 0.1 | 0.1 | 26.6 |
| | MAST | 0.262 | 1.000 | | | | | | |
| | TWO-SIGMA | **1.000** | **1.000** | | | | | | |
| 10000/10000 | ZINB | 1.000 | 1.000 | (-2, **0**, -0.5, -2) | (2, **0.03**, -0.1, 0.6) | 10 | 0.1 | 0.1 | 27.3 |
| | MAST | 0.461 | 1.000 | | | | | | |
| | TWO-SIGMA | **1.000** | **1.000** | | | | | | |
| 10000/10000 | ZINB | 1.000 | 1.000 | (-1, **0**, -0.5, -2) | (2, **0.03**, -0.1, 0.6) | 10 | 0.1 | 0.1 | 26.9 |
| | MAST | 0.246 | 1.000 | | | | | | |
| | TWO-SIGMA | **1.000** | **1.000** | | | | | | |
| 10000/10000 | ZINB | 1.000 | 1.000 | (0, **0**, -0.5, -2) | (2, **0.03**, -0.1, 0.6) | 10 | 0.1 | 0.1 | 29.8 |
| | MAST | 0.280 | 1.000 | | | | | | |
| | TWO-SIGMA | **1.000** | **1.000** | | | | | | |
| 10000/10000 | ZINB | 1.000 | 1.000 | (1, **0**, -0.5, -2) | (2, **0.03**, -0.1, 0.6) | 10 | 0.1 | 0.1 | 30.2 |
| | MAST | 0.420 | 1.000 | | | | | | |
| | TWO-SIGMA | **1.000** | **1.000** | | | | | | |
| 10000/10000 | ZINB | 1.000 | 1.000 | (2, **0**, -0.5, -2) | (2, **0.03**, -0.1, 0.6) | 10 | 0.1 | 0.1 | 29.5 |
| | MAST | 0.211 | 0.998 | | | | | | |
| | TWO-SIGMA | **0.997** | **0.997** | | | | | | |
| 10000/10000 | ZINB | 0.999 | 0.999 | (3, **0**, -0.5, -2) | (2, **0.03**, -0.1, 0.6) | 10 | 0.1 | 0.1 | 29.7 |
| | MAST | 0.134 | 0.966 | | | | | | |
| | TWO-SIGMA | **0.341** | **0.343** | | | | | | |
| 10000/10000 | ZINB | 0.495 | 0.495 | (-3, **0.03**, -0.5, -2) | (2, **0**, -0.1, 0.6) | 10 | 0.1 | 0.1 | 26.6 |
| | MAST | 0.546 | 0.368 | | | | | | |
| | TWO-SIGMA | **0.482** | **0.484** | | | | | | |
| 10000/10000 | ZINB | 0.611 | 0.611 | (-2, **0.03**, -0.5, -2) | (2, **0**, -0.1, 0.6) | 10 | 0.1 | 0.1 | 26.4 |
| | MAST | 0.703 | 0.513 | | | | | | |
| | TWO-SIGMA | **0.561** | **0.563** | | | | | | |
| 9999/10000 | ZINB | 0.668 | 0.669 | (-1, **0.03**, -0.5, -2) | (2, **0**, -0.1, 0.6) | 10 | 0.1 | 0.1 | 29.3 |
| | MAST | 0.768 | 0.588 | | | | | | |
| | TWO-SIGMA | **0.577** | **0.577** | | | | | | |
| 10000/10000 | ZINB | 0.674 | 0.676 | (0, **0.03**, -0.5, -2) | (2, **0**, -0.1, 0.6) | 10 | 0.1 | 0.1 | 30.9 |
| | MAST | 0.786 | 0.606 | | | | | | |
| | TWO-SIGMA | **0.544** | **0.546** | | | | | | |
| 10000/10000 | ZINB | 0.624 | 0.624 | (1, **0.03**, -0.5, -2) | (2, **0**, -0.1, 0.6) | 10 | 0.1 | 0.1 | 31.0 |
| | MAST | 0.768 | 0.566 | | | | | | |
| | TWO-SIGMA | **0.451** | **0.452** | | | | | | |
| 10000/10000 | ZINB | 0.526 | 0.526 | (2, **0.03**, -0.5, -2) | (2, **0**, -0.1, 0.6) | 10 | 0.1 | 0.1 | 28.6 |
| | MAST | 0.670 | 0.449 | | | | | | |
| | TWO-SIGMA | **0.343** | **0.343** | | | | | | |
| 10000/10000 | ZINB | 0.394 | 0.393 | (3, **0.03**, -0.5, -2) | (2, **0**, -0.1, 0.6) | 10 | 0.1 | 0.1 | 28.4 |
| | MAST | 0.539 | 0.318 | | | | | | |

Supplementary Table S6: Apparent Power using LRT to test $H_0 : \alpha_1 = 0, \beta_1 = 0$ with a significance level of 0.05

### Case 2: 100 individuals, 500 single-cells each, 0.05 level

##### Power Scenarios 1 & 2 from Figure 2 of main text

##### Effects in both components in either the same or different directions

| | | LRT | Combined $\chi^2$ | Simulation Parameters | | | | | Avg. Time |
|---|---|---|---|---|---|---|---|---|---|
| N / N Max | Model | P(Reject $H_0$) | P(Reject $H_0$) | $\boldsymbol{\alpha} = (\alpha_0, \alpha_1, \alpha_2, \alpha_3)$ | $\boldsymbol{\beta} = (\beta_0, \beta_1, \beta_2, \beta_3)$ | $\phi$ | $\sigma_a$ | $\sigma_b$ | (min) |
| | TWO-SIGMA | **0.849** | **0.858** | | | | | | |
| 10000/10000 | ZINB | 0.995 | 0.995 | (-3, **0.03**, -0.5, -2) | (2, **0.03**, -0.1, 0.6) | 10 | 0.1 | 0.1 | 21.6 |
| | MAST | 0.441 | 0.988 | | | | | | |
| | TWO-SIGMA | **0.868** | **0.873** | | | | | | |
| 10000/10000 | ZINB | 0.997 | 0.997 | (-2, **0.03**, -0.5, -2) | (2, **0.03**, -0.1, 0.6) | 10 | 0.1 | 0.1 | 23.9 |
| | MAST | 0.712 | 0.992 | | | | | | |
| | TWO-SIGMA | **0.884** | **0.890** | | | | | | |
| 10000/10000 | ZINB | 0.996 | 0.996 | (-1, **0.03**, -0.5, -2) | (2, **0.03**, -0.1, 0.6) | 10 | 0.1 | 0.1 | 22.6 |
| | MAST | 0.739 | 0.991 | | | | | | |
| | TWO-SIGMA | **0.888** | **0.893** | | | | | | |
| 10000/10000 | ZINB | 0.996 | 0.996 | (0, **0.03**, -0.5, -2) | (2, **0.03**, -0.1, 0.6) | 10 | 0.1 | 0.1 | 23.7 |
| | MAST | 0.784 | 0.990 | | | | | | |
| | TWO-SIGMA | **0.869** | **0.875** | | | | | | |
| 10000/10000 | ZINB | 0.993 | 0.993 | (1, **0.03**, -0.5, -2) | (2, **0.03**, -0.1, 0.6) | 10 | 0.1 | 0.1 | 26.3 |
| | MAST | 0.790 | 0.985 | | | | | | |
| | TWO-SIGMA | **0.844** | **0.850** | | | | | | |
| 10000/10000 | ZINB | 0.986 | 0.986 | (2, **0.03**, -0.5, -2) | (2, **0.03**, -0.1, 0.6) | 10 | 0.1 | 0.1 | 25.6 |
| | MAST | 0.672 | 0.964 | | | | | | |
| | TWO-SIGMA | **0.792** | **0.799** | | | | | | |
| 10000/10000 | ZINB | 0.969 | 0.969 | (3, **0.03**, -0.5, -2) | (2, **0.03**, -0.1, 0.6) | 10 | 0.1 | 0.1 | 26.1 |
| | MAST | 0.564 | 0.914 | | | | | | |
| | TWO-SIGMA | **0.853** | **0.858** | | | | | | |
| 9996/10000 | ZINB | 0.996 | 0.996 | (-3, **0.03**, -0.5, -2) | (2, **-0.03**, -0.1, 0.6) | 10 | 0.1 | 0.1 | 22.1 |
| | MAST | 0.802 | 0.991 | | | | | | |
| | TWO-SIGMA | **0.873** | **0.876** | | | | | | |
| 10000/10000 | ZINB | 0.997 | 0.997 | (-2, **0.03**, -0.5, -2) | (2, **-0.03**, -0.1, 0.6) | 10 | 0.1 | 0.1 | 23.8 |
| | MAST | 0.869 | 0.992 | | | | | | |
| | TWO-SIGMA | **0.885** | **0.889** | | | | | | |
| 9999/10000 | ZINB | 0.997 | 0.997 | (-1, **0.03**, -0.5, -2) | (2, **-0.03**, -0.1, 0.6) | 10 | 0.1 | 0.1 | 22.7 |
| | MAST | 0.839 | 0.992 | | | | | | |
| | TWO-SIGMA | **0.880** | **0.885** | | | | | | |
| 10000/10000 | ZINB | 0.996 | 0.996 | (0, **0.03**, -0.5, -2) | (2, **-0.03**, -0.1, 0.6) | 10 | 0.1 | 0.1 | 23.3 |
| | MAST | 0.839 | 0.990 | | | | | | |
| | TWO-SIGMA | **0.874** | **0.878** | | | | | | |
| 9999/10000 | ZINB | 0.993 | 0.993 | (1, **0.03**, -0.5, -2) | (2, **-0.03**, -0.1, 0.6) | 10 | 0.1 | 0.1 | 25.8 |
| | MAST | 0.824 | 0.985 | | | | | | |
| | TWO-SIGMA | **0.840** | **0.846** | | | | | | |
| 10000/10000 | ZINB | 0.989 | 0.989 | (2, **0.03**, -0.5, -2) | (2, **-0.03**, -0.1, 0.6) | 10 | 0.1 | 0.1 | 25.9 |
| | MAST | 0.709 | 0.965 | | | | | | |
| | TWO-SIGMA | **0.799** | **0.806** | | | | | | |
| 10000/10000 | ZINB | 0.974 | 0.974 | (3, **0.03**, -0.5, -2) | (2, **-0.03**, -0.1, 0.6) | 10 | 0.1 | 0.1 | 25.6 |
| | MAST | 0.573 | 0.919 | | | | | | |

Supplementary Table S7: Apparent Power using LRT to test $H_0 : \alpha_1 = 0, \beta_1 = 0$ with a significance level of 0.05

### Case 2: 100 individuals, 500 single-cells each, 0.05 level

##### Power Scenarios 3 & 4 from Figure 2 of main text

##### Effects in one component at a time

| | | LRT | Combined $\chi^2$ | Simulation Parameters | | | | | Avg. Time |
|---|---|---|---|---|---|---|---|---|---|
| N / N Max | Model | P(Reject $H_0$) | P(Reject $H_0$) | $\boldsymbol{\alpha} = (\alpha_0, \alpha_1, \alpha_2, \alpha_3)$ | $\boldsymbol{\beta} = (\beta_0, \beta_1, \beta_2, \beta_3)$ | $\phi$ | $\sigma_a$ | $\sigma_b$ | (min) |
| 9999/10000 | TWO-SIGMA | **0.748** | **0.757** | (-3, **0**, -0.5, -2) | (2, **0.03**, -0.1, 0.6) | 10 | 0.1 | 0.1 | 21.9 |
| | ZINB | 0.992 | 0.992 | | | | | | |
| | MAST | 0.341 | 0.982 | | | | | | |
| 10000/10000 | TWO-SIGMA | **0.744** | **0.754** | (-2, **0**, -0.5, -2) | (2, **0.03**, -0.1, 0.6) | 10 | 0.1 | 0.1 | 22.7 |
| | ZINB | 0.993 | 0.993 | | | | | | |
| | MAST | 0.478 | 0.981 | | | | | | |
| 10000/10000 | TWO-SIGMA | **0.734** | **0.744** | (-1, **0**, -0.5, -2) | (2, **0.03**, -0.1, 0.6) | 10 | 0.1 | 0.1 | 21.8 |
| | ZINB | 0.991 | 0.991 | | | | | | |
| | MAST | 0.347 | 0.978 | | | | | | |
| 10000/10000 | TWO-SIGMA | **0.738** | **0.746** | (0, **0**, -0.5, -2) | (2, **0.03**, -0.1, 0.6) | 10 | 0.1 | 0.1 | 22.1 |
| | ZINB | 0.991 | 0.990 | | | | | | |
| | MAST | 0.369 | 0.975 | | | | | | |
| 10000/10000 | TWO-SIGMA | **0.725** | **0.735** | (1, **0**, -0.5, -2) | (2, **0.03**, -0.1, 0.6) | 10 | 0.1 | 0.1 | 25.4 |
| | ZINB | 0.984 | 0.984 | | | | | | |
| | MAST | 0.444 | 0.962 | | | | | | |
| 10000/10000 | TWO-SIGMA | **0.699** | **0.711** | (2, **0**, -0.5, -2) | (2, **0.03**, -0.1, 0.6) | 10 | 0.1 | 0.1 | 24.9 |
| | ZINB | 0.974 | 0.974 | | | | | | |
| | MAST | 0.282 | 0.923 | | | | | | |
| 10000/10000 | TWO-SIGMA | **0.664** | **0.677** | (3, **0**, -0.5, -2) | (2, **0.03**, -0.1, 0.6) | 10 | 0.1 | 0.1 | 26.6 |
| | ZINB | 0.949 | 0.949 | | | | | | |
| | MAST | 0.187 | 0.851 | | | | | | |
| 9999/10000 | TWO-SIGMA | **0.279** | **0.289** | (-3, **0.03**, -0.5, -2) | (2, **0**, -0.1, 0.6) | 10 | 0.1 | 0.1 | 22.7 |
| | ZINB | 0.770 | 0.769 | | | | | | |
| | MAST | 0.558 | 0.654 | | | | | | |
| 9999/10000 | TWO-SIGMA | **0.355** | **0.367** | (-2, **0.03**, -0.5, -2) | (2, **0**, -0.1, 0.6) | 10 | 0.1 | 0.1 | 21.1 |
| | ZINB | 0.819 | 0.820 | | | | | | |
| | MAST | 0.688 | 0.729 | | | | | | |
| 10000/10000 | TWO-SIGMA | **0.388** | **0.400** | (-1, **0.03**, -0.5, -2) | (2, **0**, -0.1, 0.6) | 10 | 0.1 | 0.1 | 19.7 |
| | ZINB | 0.834 | 0.834 | | | | | | |
| | MAST | 0.732 | 0.756 | | | | | | |
| 10000/10000 | TWO-SIGMA | **0.398** | **0.408** | (0, **0.03**, -0.5, -2) | (2, **0**, -0.1, 0.6) | 10 | 0.1 | 0.1 | 20.6 |
| | ZINB | 0.823 | 0.823 | | | | | | |
| | MAST | 0.755 | 0.744 | | | | | | |
| 9899/10000 | TWO-SIGMA | **0.380** | **0.391** | (1, **0.03**, -0.5, -2) | (2, **0**, -0.1, 0.6) | 10 | 0.1 | 0.1 | 25.3 |
| | ZINB | 0.784 | 0.784 | | | | | | |
| | MAST | 0.754 | 0.706 | | | | | | |
| 10000/10000 | TWO-SIGMA | **0.339** | **0.349** | (2, **0.03**, -0.5, -2) | (2, **0**, -0.1, 0.6) | 10 | 0.1 | 0.1 | 26.4 |
| | ZINB | 0.718 | 0.719 | | | | | | |
| | MAST | 0.651 | 0.601 | | | | | | |
| 10000/10000 | TWO-SIGMA | **0.270** | **0.280** | (3, **0.03**, -0.5, -2) | (2, **0**, -0.1, 0.6) | 10 | 0.1 | 0.1 | 26.1 |
| | ZINB | 0.588 | 0.588 | | | | | | |
| | MAST | 0.533 | 0.446 | | | | | | |

Supplementary Table S8: Apparent Power using LRT to test $H_0 : \alpha_1 = 0, \beta_1 = 0$ with a significance level of 0.05

#### 4.2 Results using "True" Power for MAST and the ZINB model

### Case 1: 1000 individuals, 50 single-cells each, 0.05 level

##### Power Scenarios 1 & 2 from Figure 2 of main text
##### Effects in both components in either the same or different directions

| | | LRT | Combined $\chi^2$ | Simulation Parameters | | | | | Avg. Time |
|---|---|---|---|---|---|---|---|---|---|
| N / N Max | Model | P(Reject $H_0$) | P(Reject $H_0$) | $\boldsymbol{\alpha} = (\alpha_0, \alpha_1, \alpha_2, \alpha_3)$ | $\boldsymbol{\beta} = (\beta_0, \beta_1, \beta_2, \beta_3)$ | $\phi$ | $\sigma_a$ | $\sigma_b$ | (min) |
| | TWO-SIGMA | **1.000** | **1.000** | | | | | | |
| 10000/10000 | ZINB | 1.000 | 1.000 | (-3, **0.03**, -0.5, -2) | (2, **0.03**, -0.1, 0.6) | 10 | 0.1 | 0.1 | 27.9 |
| | MAST | 0.194 | 0.352 | | | | | | |
| | TWO-SIGMA | **1.000** | **1.000** | | | | | | |
| 10000/10000 | ZINB | 1.000 | 1.000 | (-2, **0.03**, -0.5, -2) | (2, **0.03**, -0.1, 0.6) | 10 | 0.1 | 0.1 | 27.8 |
| | MAST | 0.588 | 0.721 | | | | | | |
| | TWO-SIGMA | **1.000** | **1.000** | | | | | | |
| 10000/10000 | ZINB | 1.000 | 1.000 | (-1, **0.03**, -0.5, -2) | (2, **0.03**, -0.1, 0.6) | 10 | 0.1 | 0.1 | 27.0 |
| | MAST | 0.569 | 0.746 | | | | | | |
| | TWO-SIGMA | **1.000** | **1.000** | | | | | | |
| 10000/10000 | ZINB | 1.000 | 1.000 | (0, **0.03**, -0.5, -2) | (2, **0.03**, -0.1, 0.6) | 10 | 0.1 | 0.1 | 29.4 |
| | MAST | 0.644 | 0.802 | | | | | | |
| | TWO-SIGMA | **1.000** | **1.000** | | | | | | |
| 10000/10000 | ZINB | 1.000 | 1.000 | (1, **0.03**, -0.5, -2) | (2, **0.03**, -0.1, 0.6) | 10 | 0.1 | 0.1 | 29.4 |
| | MAST | 0.705 | 0.822 | | | | | | |
| | TWO-SIGMA | **1.000** | **1.000** | | | | | | |
| 10000/10000 | ZINB | 1.000 | 1.000 | (2, **0.03**, -0.5, -2) | (2, **0.03**, -0.1, 0.6) | 10 | 0.1 | 0.1 | 27.5 |
| | MAST | 0.524 | 0.700 | | | | | | |
| | TWO-SIGMA | **0.999** | **0.999** | | | | | | |
| 10000/10000 | ZINB | 0.998 | 0.998 | (3, **0.03**, -0.5, -2) | (2, **0.03**, -0.1, 0.6) | 10 | 0.1 | 0.1 | 27.2 |
| | MAST | 0.358 | 0.547 | | | | | | |
| | TWO-SIGMA | **1.000** | **1.000** | | | | | | |
| 10000/10000 | ZINB | 1.000 | 1.000 | (-3, **0.03**, -0.5, -2) | (2, **-0.03**, -0.1, 0.6) | 10 | 0.1 | 0.1 | 27.4 |
| | MAST | 0.673 | 0.829 | | | | | | |
| | TWO-SIGMA | **1.000** | **1.000** | | | | | | |
| 10000/10000 | ZINB | 1.000 | 1.000 | (-2, **0.03**, -0.5, -2) | (2, **-0.03**, -0.1, 0.6) | 10 | 0.1 | 0.1 | 27.6 |
| | MAST | 0.803 | 0.904 | | | | | | |
| | TWO-SIGMA | **1.000** | **1.000** | | | | | | |
| 10000/10000 | ZINB | 1.000 | 1.000 | (-1, **0.03**, -0.5, -2) | (2, **-0.03**, -0.1, 0.6) | 10 | 0.1 | 0.1 | 27.7 |
| | MAST | 0.736 | 0.872 | | | | | | |
| | TWO-SIGMA | **1.000** | **1.000** | | | | | | |
| 10000/10000 | ZINB | 1.000 | 1.000 | (0, **0.03**, -0.5, -2) | (2, **-0.03**, -0.1, 0.6) | 10 | 0.1 | 0.1 | 29.5 |
| | MAST | 0.741 | 0.867 | | | | | | |
| | TWO-SIGMA | **1.000** | **1.000** | | | | | | |
| 10000/10000 | ZINB | 1.000 | 1.000 | (1, **0.03**, -0.5, -2) | (2, **-0.03**, -0.1, 0.6) | 10 | 0.1 | 0.1 | 28.2 |
| | MAST | 0.738 | 0.852 | | | | | | |
| | TWO-SIGMA | **1.000** | **1.000** | | | | | | |
| 10000/10000 | ZINB | 1.000 | 1.000 | (2, **0.03**, -0.5, -2) | (2, **-0.03**, -0.1, 0.6) | 10 | 0.1 | 0.1 | 27.0 |
| | MAST | 0.541 | 0.715 | | | | | | |
| | TWO-SIGMA | **0.998** | **0.998** | | | | | | |
| 10000/10000 | ZINB | 0.997 | 0.997 | (3, **0.03**, -0.5, -2) | (2, **-0.03**, -0.1, 0.6) | 10 | 0.1 | 0.1 | 26.2 |
| | MAST | 0.369 | 0.557 | | | | | | |

Supplementary Table S9: True Power using LRT to test $H_0 : \alpha_1 = 0,\ \beta_1 = 0$ with a significance level of 0.05

### Case 1: 1000 individuals, 50 single-cells each, 0.05 level

##### Power Scenarios 3 & 4 from Figure 2 of main text

##### Effects in one component at a time

| N / N Max | Model | LRT P(Reject $H_0$) | Combined $\chi^2$ P(Reject $H_0$) | Simulation Parameters $\boldsymbol{\alpha} = (\alpha_0, \alpha_1, \alpha_2, \alpha_3)$ | $\boldsymbol{\beta} = (\beta_0, \beta_1, \beta_2, \beta_3)$ | $\phi$ | $\sigma_a$ | $\sigma_b$ | Avg. Time (min) |
|---|---|---|---|---|---|---|---|---|---|
| | TWO-SIGMA | **1.000** | **1.000** | | | | | | |
| 10000/10000 | ZINB | 1.000 | 1.000 | (-3, **0**, -0.5, -2) | (2, **0.03**, -0.1, 0.6) | 10 | 0.1 | 0.1 | 26.6 |
| | MAST | 0.119 | 0.242 | | | | | | |
| | TWO-SIGMA | **1.000** | **1.000** | | | | | | |
| 10000/10000 | ZINB | 1.000 | 1.000 | (-2, **0**, -0.5, -2) | (2, **0.03**, -0.1, 0.6) | 10 | 0.1 | 0.1 | 27.3 |
| | MAST | 0.366 | 0.450 | | | | | | |
| | TWO-SIGMA | **1.000** | **1.000** | | | | | | |
| 10000/10000 | ZINB | 1.000 | 1.000 | (-1, **0**, -0.5, -2) | (2, **0.03**, -0.1, 0.6) | 10 | 0.1 | 0.1 | 26.9 |
| | MAST | 0.116 | 0.229 | | | | | | |
| | TWO-SIGMA | **1.000** | **1.000** | | | | | | |
| 10000/10000 | ZINB | 1.000 | 1.000 | (0, **0**, -0.5, -2) | (2, **0.03**, -0.1, 0.6) | 10 | 0.1 | 0.1 | 29.8 |
| | MAST | 0.168 | 0.263 | | | | | | |
| | TWO-SIGMA | **1.000** | **1.000** | | | | | | |
| 10000/10000 | ZINB | 1.000 | 1.000 | (1, **0**, -0.5, -2) | (2, **0.03**, -0.1, 0.6) | 10 | 0.1 | 0.1 | 30.2 |
| | MAST | 0.361 | 0.411 | | | | | | |
| | TWO-SIGMA | **1.000** | **1.000** | | | | | | |
| 10000/10000 | ZINB | 1.000 | 1.000 | (2, **0**, -0.5, -2) | (2, **0.03**, -0.1, 0.6) | 10 | 0.1 | 0.1 | 29.5 |
| | MAST | 0.124 | 0.200 | | | | | | |
| | TWO-SIGMA | **0.997** | **0.997** | | | | | | |
| 10000/10000 | ZINB | 0.996 | 0.996 | (3, **0**, -0.5, -2) | (2, **0.03**, -0.1, 0.6) | 10 | 0.1 | 0.1 | 29.7 |
| | MAST | 0.048 | 0.120 | | | | | | |
| | TWO-SIGMA | **0.336** | **0.335** | | | | | | |
| 10000/10000 | ZINB | 0.295 | 0.294 | (-3, **0.03**, -0.5, -2) | (2, **0**, -0.1, 0.6) | 10 | 0.1 | 0.1 | 26.6 |
| | MAST | 0.329 | 0.522 | | | | | | |
| | TWO-SIGMA | **0.476** | **0.474** | | | | | | |
| 10000/10000 | ZINB | 0.401 | 0.400 | (-2, **0.03**, -0.5, -2) | (2, **0**, -0.1, 0.6) | 10 | 0.1 | 0.1 | 26.4 |
| | MAST | 0.489 | 0.684 | | | | | | |
| | TWO-SIGMA | **0.554** | **0.553** | | | | | | |
| 9999/10000 | ZINB | 0.459 | 0.459 | (-1, **0.03**, -0.5, -2) | (2, **0**, -0.1, 0.6) | 10 | 0.1 | 0.1 | 29.3 |
| | MAST | 0.573 | 0.747 | | | | | | |
| | TWO-SIGMA | **0.570** | **0.568** | | | | | | |
| 10000/10000 | ZINB | 0.465 | 0.464 | (0, **0.03**, -0.5, -2) | (2, **0**, -0.1, 0.6) | 10 | 0.1 | 0.1 | 30.9 |
| | MAST | 0.601 | 0.769 | | | | | | |
| | TWO-SIGMA | **0.539** | **0.537** | | | | | | |
| 10000/10000 | ZINB | 0.409 | 0.408 | (1, **0.03**, -0.5, -2) | (2, **0**, -0.1, 0.6) | 10 | 0.1 | 0.1 | 31.0 |
| | MAST | 0.579 | 0.750 | | | | | | |
| | TWO-SIGMA | **0.445** | **0.444** | | | | | | |
| 10000/10000 | ZINB | 0.306 | 0.305 | (2, **0.03**, -0.5, -2) | (2, **0**, -0.1, 0.6) | 10 | 0.1 | 0.1 | 28.6 |
| | MAST | 0.459 | 0.649 | | | | | | |
| | TWO-SIGMA | **0.337** | **0.335** | | | | | | |
| 10000/10000 | ZINB | 0.205 | 0.204 | (3, **0.03**, -0.5, -2) | (2, **0**, -0.1, 0.6) | 10 | 0.1 | 0.1 | 28.4 |
| | MAST | 0.331 | 0.518 | | | | | | |

Supplementary Table S10: True Power using LRT to test $H_0 : \alpha_1 = 0,\ \beta_1 = 0$ with a significance level of 0.05

### Case 2: 100 individuals, 500 single-cells each, 0.05 level

##### Power Scenarios 1 & 2 from Figure 2 of main text

##### Effects in both components in either the same or different directions

| N / N Max | Model | LRT P(Reject $H_0$) | Combined $\chi^2$ P(Reject $H_0$) | Simulation Parameters $\boldsymbol{\alpha} = (\alpha_0, \alpha_1, \alpha_2, \alpha_3)$ | $\boldsymbol{\beta} = (\beta_0, \beta_1, \beta_2, \beta_3)$ | $\phi$ | $\sigma_a$ | $\sigma_b$ | Avg. Time (min) |
|---|---|---|---|---|---|---|---|---|---|
| | TWO-SIGMA | **0.837** | **0.830** | | | | | | |
| 10000/10000 | ZINB | 0.937 | 0.937 | (-3, **0.03**, -0.5, -2) | (2, **0.03**, -0.1, 0.6) | 10 | 0.1 | 0.1 | 21.6 |
| | MAST | 0.052 | 0.085 | | | | | | |
| | TWO-SIGMA | **0.858** | **0.852** | | | | | | |
| 10000/10000 | ZINB | 0.935 | 0.934 | (-2, **0.03**, -0.5, -2) | (2, **0.03**, -0.1, 0.6) | 10 | 0.1 | 0.1 | 23.9 |
| | MAST | 0.215 | 0.298 | | | | | | |
| | TWO-SIGMA | **0.872** | **0.866** | | | | | | |
| 10000/10000 | ZINB | 0.926 | 0.926 | (-1, **0.03**, -0.5, -2) | (2, **0.03**, -0.1, 0.6) | 10 | 0.1 | 0.1 | 22.6 |
| | MAST | 0.207 | 0.290 | | | | | | |
| | TWO-SIGMA | **0.876** | **0.870** | | | | | | |
| 10000/10000 | ZINB | 0.899 | 0.899 | (0, **0.03**, -0.5, -2) | (2, **0.03**, -0.1, 0.6) | 10 | 0.1 | 0.1 | 23.7 |
| | MAST | 0.298 | 0.373 | | | | | | |
| | TWO-SIGMA | **0.859** | **0.852** | | | | | | |
| 10000/10000 | ZINB | 0.840 | 0.840 | (1, **0.03**, -0.5, -2) | (2, **0.03**, -0.1, 0.6) | 10 | 0.1 | 0.1 | 26.3 |
| | MAST | 0.394 | 0.456 | | | | | | |
| | TWO-SIGMA | **0.831** | **0.822** | | | | | | |
| 10000/10000 | ZINB | 0.729 | 0.728 | (2, **0.03**, -0.5, -2) | (2, **0.03**, -0.1, 0.6) | 10 | 0.1 | 0.1 | 25.6 |
| | MAST | 0.191 | 0.252 | | | | | | |
| | TWO-SIGMA | **0.778** | **0.768** | | | | | | |
| 10000/10000 | ZINB | 0.502 | 0.500 | (3, **0.03**, -0.5, -2) | (2, **0.03**, -0.1, 0.6) | 10 | 0.1 | 0.1 | 26.1 |
| | MAST | 0.081 | 0.130 | | | | | | |
| | TWO-SIGMA | **0.842** | **0.835** | | | | | | |
| 9996/10000 | ZINB | 0.943 | 0.943 | (-3, **0.03**, -0.5, -2) | (2, **-0.03**, -0.1, 0.6) | 10 | 0.1 | 0.1 | 22.1 |
| | MAST | 0.257 | 0.350 | | | | | | |
| | TWO-SIGMA | **0.862** | **0.855** | | | | | | |
| 10000/10000 | ZINB | 0.937 | 0.937 | (-2, **0.03**, -0.5, -2) | (2, **-0.03**, -0.1, 0.6) | 10 | 0.1 | 0.1 | 23.8 |
| | MAST | 0.388 | 0.483 | | | | | | |
| | TWO-SIGMA | **0.875** | **0.868** | | | | | | |
| 9999/10000 | ZINB | 0.927 | 0.926 | (-1, **0.03**, -0.5, -2) | (2, **-0.03**, -0.1, 0.6) | 10 | 0.1 | 0.1 | 22.7 |
| | MAST | 0.323 | 0.417 | | | | | | |
| | TWO-SIGMA | **0.870** | **0.864** | | | | | | |
| 10000/10000 | ZINB | 0.901 | 0.900 | (0, **0.03**, -0.5, -2) | (2, **-0.03**, -0.1, 0.6) | 10 | 0.1 | 0.1 | 23.3 |
| | MAST | 0.371 | 0.452 | | | | | | |
| | TWO-SIGMA | **0.862** | **0.855** | | | | | | |
| 9999/10000 | ZINB | 0.847 | 0.846 | (1, **0.03**, -0.5, -2) | (2, **-0.03**, -0.1, 0.6) | 10 | 0.1 | 0.1 | 25.8 |
| | MAST | 0.434 | 0.503 | | | | | | |
| | TWO-SIGMA | **0.827** | **0.819** | | | | | | |
| 10000/10000 | ZINB | 0.717 | 0.716 | (2, **0.03**, -0.5, -2) | (2, **-0.03**, -0.1, 0.6) | 10 | 0.1 | 0.1 | 25.9 |
| | MAST | 0.206 | 0.278 | | | | | | |
| | TWO-SIGMA | **0.785** | **0.775** | | | | | | |
| 10000/10000 | ZINB | 0.503 | 0.500 | (3, **0.03**, -0.5, -2) | (2, **-0.03**, -0.1, 0.6) | 10 | 0.1 | 0.1 | 25.6 |
| | MAST | 0.079 | 0.132 | | | | | | |

Supplementary Table S11: True Power using LRT to test $H_0 : \alpha_1 = 0,\ \beta_1 = 0$ with a significance level of 0.05

### Case 2: 100 individuals, 500 single-cells each, 0.05 level

#### Power Scenarios 3 & 4 from Figure 2 of main text

#### Effects in one component at a time

| N / N Max | Model | LRT P(Reject $H_0$) | Combined $\chi^2$ P(Reject $H_0$) | Simulation Parameters $\boldsymbol{\alpha} = (\alpha_0, \alpha_1, \alpha_2, \alpha_3)$ | $\boldsymbol{\beta} = (\beta_0, \beta_1, \beta_2, \beta_3)$ | $\phi$ | $\sigma_a$ | $\sigma_b$ | Avg. Time (min) |
|---|---|---|---|---|---|---|---|---|---|
| 9999/10000 | TWO-SIGMA | **0.730** | **0.719** | (-3, **0**, -0.5, -2) | (2, **0.03**, -0.1, 0.6) | 10 | 0.1 | 0.1 | 21.9 |
| | ZINB | 0.932 | 0.932 | | | | | | |
| | MAST | 0.037 | 0.058 | | | | | | |
| 10000/10000 | TWO-SIGMA | **0.725** | **0.716** | (-2, **0**, -0.5, -2) | (2, **0.03**, -0.1, 0.6) | 10 | 0.1 | 0.1 | 22.7 |
| | ZINB | 0.925 | 0.924 | | | | | | |
| | MAST | 0.109 | 0.152 | | | | | | |
| 10000/10000 | TWO-SIGMA | **0.715** | **0.704** | (-1, **0**, -0.5, -2) | (2, **0.03**, -0.1, 0.6) | 10 | 0.1 | 0.1 | 21.8 |
| | ZINB | 0.907 | 0.906 | | | | | | |
| | MAST | 0.038 | 0.059 | | | | | | |
| 10000/10000 | TWO-SIGMA | **0.723** | **0.713** | (0, **0**, -0.5, -2) | (2, **0.03**, -0.1, 0.6) | 10 | 0.1 | 0.1 | 22.1 |
| | ZINB | 0.870 | 0.870 | | | | | | |
| | MAST | 0.128 | 0.142 | | | | | | |
| 10000/10000 | TWO-SIGMA | **0.710** | **0.699** | (1, **0**, -0.5, -2) | (2, **0.03**, -0.1, 0.6) | 10 | 0.1 | 0.1 | 25.4 |
| | ZINB | 0.810 | 0.809 | | | | | | |
| | MAST | 0.265 | 0.278 | | | | | | |
| 10000/10000 | TWO-SIGMA | **0.680** | **0.669** | (2, **0**, -0.5, -2) | (2, **0.03**, -0.1, 0.6) | 10 | 0.1 | 0.1 | 24.9 |
| | ZINB | 0.675 | 0.673 | | | | | | |
| | MAST | 0.070 | 0.083 | | | | | | |
| 10000/10000 | TWO-SIGMA | **0.645** | **0.632** | (3, **0**, -0.5, -2) | (2, **0.03**, -0.1, 0.6) | 10 | 0.1 | 0.1 | 26.6 |
| | ZINB | 0.452 | 0.451 | | | | | | |
| | MAST | 0.012 | 0.020 | | | | | | |
| 9999/10000 | TWO-SIGMA | **0.263** | **0.252** | (-3, **0.03**, -0.5, -2) | (2, **0**, -0.1, 0.6) | 10 | 0.1 | 0.1 | 22.7 |
| | ZINB | 0.184 | 0.184 | | | | | | |
| | MAST | 0.081 | 0.129 | | | | | | |
| 9999/10000 | TWO-SIGMA | **0.337** | **0.324** | (-2, **0.03**, -0.5, -2) | (2, **0**, -0.1, 0.6) | 10 | 0.1 | 0.1 | 21.1 |
| | ZINB | 0.184 | 0.184 | | | | | | |
| | MAST | 0.161 | 0.238 | | | | | | |
| 10000/10000 | TWO-SIGMA | **0.368** | **0.356** | (-1, **0.03**, -0.5, -2) | (2, **0**, -0.1, 0.6) | 10 | 0.1 | 0.1 | 19.7 |
| | ZINB | 0.162 | 0.160 | | | | | | |
| | MAST | 0.206 | 0.290 | | | | | | |
| 10000/10000 | TWO-SIGMA | **0.375** | **0.364** | (0, **0.03**, -0.5, -2) | (2, **0**, -0.1, 0.6) | 10 | 0.1 | 0.1 | 20.6 |
| | ZINB | 0.128 | 0.126 | | | | | | |
| | MAST | 0.250 | 0.335 | | | | | | |
| 9899/10000 | TWO-SIGMA | **0.358** | **0.346** | (1, **0.03**, -0.5, -2) | (2, **0**, -0.1, 0.6) | 10 | 0.1 | 0.1 | 25.3 |
| | ZINB | 0.086 | 0.085 | | | | | | |
| | MAST | 0.251 | 0.336 | | | | | | |
| 10000/10000 | TWO-SIGMA | **0.321** | **0.310** | (2, **0.03**, -0.5, -2) | (2, **0**, -0.1, 0.6) | 10 | 0.1 | 0.1 | 26.4 |
| | ZINB | 0.042 | 0.042 | | | | | | |
| | MAST | 0.136 | 0.206 | | | | | | |
| 10000/10000 | TWO-SIGMA | **0.253** | **0.242** | (3, **0.03**, -0.5, -2) | (2, **0**, -0.1, 0.6) | 10 | 0.1 | 0.1 | 26.1 |
| | ZINB | 0.011 | 0.011 | | | | | | |
| | MAST | 0.068 | 0.115 | | | | | | |

Supplementary Table S12: True Power using LRT to test $H_0 : \alpha_1 = 0, \beta_1 = 0$ with a significance level of 0.05

### 5 Additional Computational Details

Suppose we have $n$ samples, each with $n_i$ single cells, $i = 1, ..., n$, the marginal likelihood $\mathrm{L}(\boldsymbol{\alpha}, \boldsymbol{\beta}, \phi, \sigma_a^2, \sigma_b^2)$ equation for the TWO-SIGMA model is given by

$$\prod_{i=1}^{n} \int_{-\infty}^{\infty} \int_{-\infty}^{\infty} \prod_{j=1}^{n_i} \Big( [\mathrm{P}(Y_{ij} = 0)]^{\mathrm{I}(y_{ij}=0)} [\mathrm{P}(Y_{ij} = y_{ij})]^{\mathrm{I}(y_{ij}>0)} \times g(a_i, b_i \mid \sigma_a^2, \sigma_b^2) \mathrm{d}a_i \mathrm{d}b_i \Big)$$

where $g(a_i, b_i \mid \sigma_a^2, \sigma_b^2)$ is the product of two normal densities (assuming $a_i \perp\!\!\!\perp b_i$).
Because no analytic solutions to this integral are available, the marginal likelihood must be approximated to obtain parameter estimates. Models that include many random effects can be fit efficiently using the implementation in the **twosigma** R package because the Laplace approximation is used to integrate out random effects and automatic differentiation is used to compute gradients [4]. It can be shown that the Laplace approximation is equivalent to using Gaussian quadrature with one quadrature point [5]. Although estimates can be biased, the Laplace approximation often performs well for count response variables [6]. For further comments on situations in which the Laplace approximation performs suitably well in practical applications, including the analysis of count data, see [7]. Finally, others have demonstrated that the Laplace approximation works quite well in non-linear mixed-effects models [8]. This framework also does not require balanced data, as is sometimes assumed for mixed-effects models; for instance, balanced data are implicitly included in the setup of [9].

### 6 More Discussion of the Zero-Inflation Component

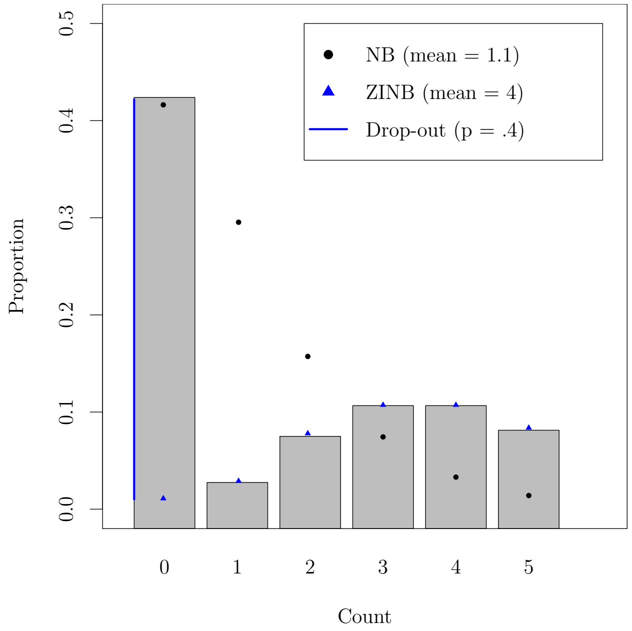


Supplementary Figure S9: Failing to account for zero-inflation: shows the impact of failing to account for zero-inflation on mean inference. Data was simulated according to a zero-inflated negative binomial (ZINB) distribution with mean 4 and drop-out probability 0.4 as specified in equation (1) in the main text. Black dots represent an approximate negative binomial fit to the data and blue represents the truth as simulated from a ZINB distribution.

Figure S9 provides a simple illustration of how ignoring this zero-inflation can lead to substantial underestimation of mean parameters. Our experience suggests that scRNA-seq data often benefit from, if not require, such explicit modelling of excess zeros. Some genes may not require the full TWO-SIGMA specification. For example, consider the marker genes of each cell type in the pancreas dataset—*GCG* for alpha cells and *INS* for beta cells [10]. These marker genes are not indicative of a need for zero inflation—only 1 alpha cell has a read count of zero for *GCG* and all beta cells have non-zero read counts for *INS*. In both cases one can remove the entire zero-inflation component and refit, continuing to allow for overdispersion and within-sample correlation. As an aside, TWO-SIGMA can estimate the zero-inflated coefficients $\boldsymbol{\alpha}$ for these genes because information about $\boldsymbol{\alpha}$ is contained in the second line of equation (1) of the main text. Doing so would seem inappropriate, however, given the inconsistency between the observed data and the idea that excess zeros are present.
